# Myosin 10 and a Cytoneme-Localized Ligand Complex Promote Morphogen Transport

**DOI:** 10.1101/2020.05.06.080820

**Authors:** Eric T. Hall, Daniel P. Stewart, Miriam Dillard, Ben Wagner, April Sykes, Jamshid Temirov, Richard E. Cheney, Motomi Mori, Camenzind G. Robinson, Stacey K. Ogden

## Abstract

Morphogens function in concentration-dependent manners to instruct cell fate during tissue patterning. Molecular mechanisms by which these signaling gradients are established and reinforced remain enigmatic. The cytoneme transport model posits that specialized filopodia extend between morphogen-sending and responding cells to ensure that appropriate signal activation thresholds are achieved across developing tissues. How morphogens are transported along and deployed from cytonemes is not known. Herein we show that the actin motor Myosin 10 promotes cytoneme-based transport of Sonic Hedgehog (SHH) morphogen to filopodial tips, and that SHH movement within cytonemes occurs by vesicular transport. We demonstrate that cytoneme-mediated deposition of SHH onto receiving cells induces a rapid signal response, and that SHH cytonemes are promoted by a complex containing a ligand-specific deployment protein and associated co-receptor.

**One-Sentence summary:** Cytoneme-based delivery of the Sonic Hedgehog activation signal is promoted by Myosin 10 and BOC/CDON co-receptor function.

## Results and Discussion

### SHH promotes cytoneme formation

Previous studies demonstrated that Hedgehog (HH) family ligands are transported along cytonemes, and that HH pathway components including the deployment protein Dispatched (DISP), the HH receptor Patched (PTCH), and co-receptors CDON/iHOG and BOC/BOI localize to the specialized filopodia (*1*–*5*). However, cell biological processes controlling cytoneme biogenesis, function, and protein entry into the structures are not yet clear. To begin to interrogate these activities, NIH3T3 cells and mouse embryonic fibroblasts (MEFs) expressing SHH plus the membrane marker mCherry-Mem (mCherry fused to the first twenty residues of neuromodulin) were fixed using the MEM-fix technique, and then examined by confocal microscopy (Figure 1A-B’, S1) (*1*, *6*). Image analysis of the mCherry-Mem signal in SHH-expressing cells revealed long extensions from both NIH3T3 cells and MEFs that reached around and over neighboring cells (Figures 1A-B, S1). Depth analysis of the mCherry-Mem signal revealed that cytoneme-like projections originated from cell membranes not in contact with the growth surface (Figure 1A’, B’, S1A-B). The small diameter, long lengths, and growth patterns of these extensions are consistent with the documented characteristics of cytonemes, indicating NIH3T3 cells and MEFs can be used to interrogate the specialized filopodia (*1*, *6*–*8*).

**Figure 1:**
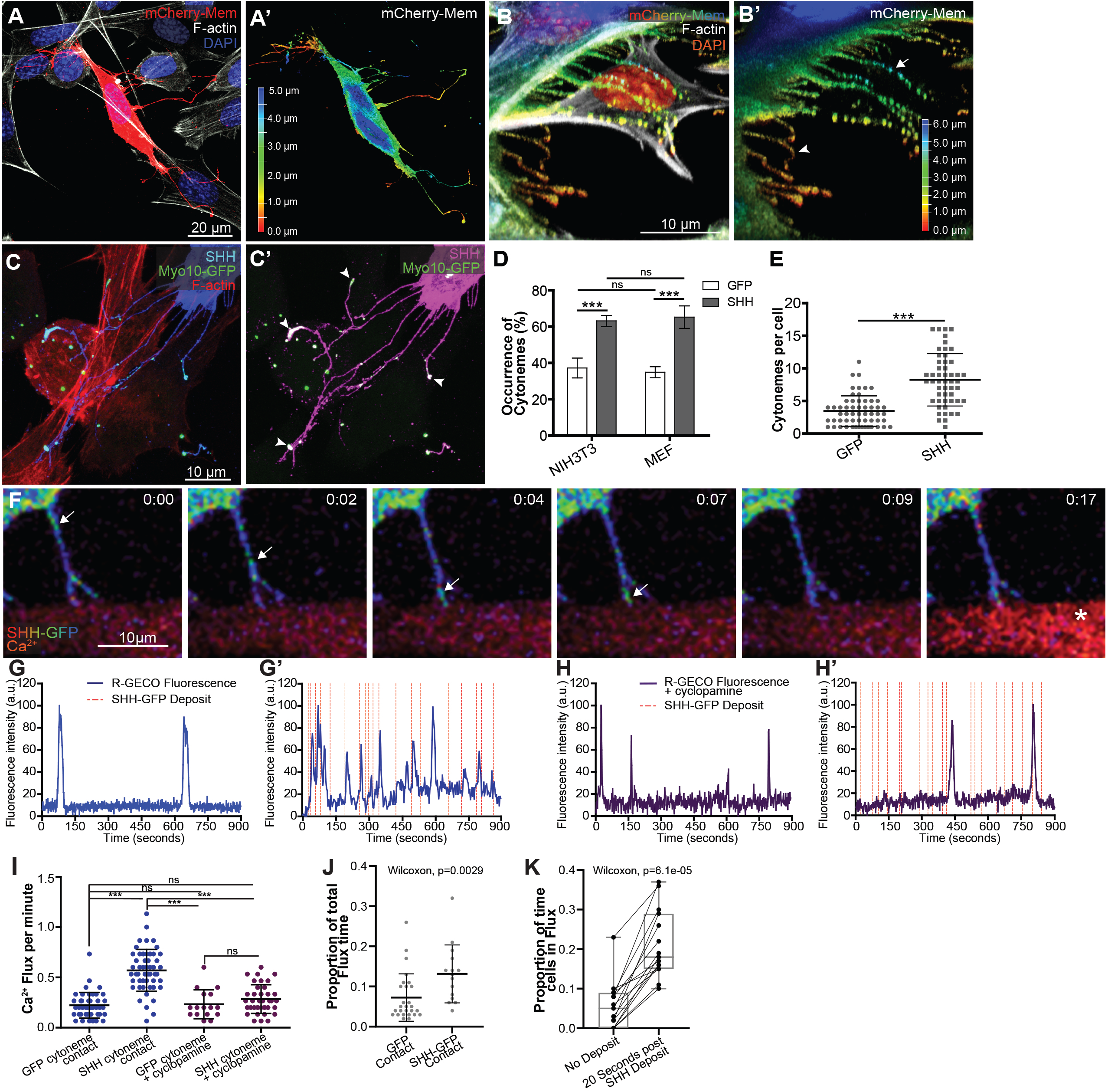
SHH promotes cytoneme formation for signal delivery. (**A-A’**) Cytonemes of a mCherry-Mem expressing cell (red) contact neighboring cells. Z-axis depth colored projection of the mCherry-Mem signal is shown in **A’**. (**B**) A 3D render of a MEF cell expressing SHH and mCherry-Mem colored for Z-axis depth shows cytonemes reaching around a neighboring cell (green, arrow in B’). Basal projecting filopodia are red, with an example indicated by an arrowhead in **B’**. (**C**) An NIH3T3 cell expressing MYO10-GFP and SHH (blue) is shown. Actin is red. SHH signal is saturated in **C’** (magenta) to visualize cytonemes. MYO10-GFP accumulates in puncta at cytoneme tips with SHH (**C’** arrow heads). (**D**) Cytoneme occurrence rates were calculated in MEM-fixed NIH3T3 and MEF cells in the absence and presence of SHH. SHH expression increases cytoneme occurrence. (**E**) The number of cytonemes per NIH3T3 cell were counted in the presence of either GFP (n=57) or SHH expression (n=51). (**F**) Time lapse images of a SHH-expressing NIH3T3 cell showing SHH-GFP as a fluorescent intensity-spectrum in puncta. Progressive movement of a puncta traveling down a cytoneme to the tip in contact with a R-GECO sensor cell is indicated by an arrow. R-GECO fluorescent intensity increases ~10 seconds after release of the SHH puncta from the cytoneme (asterisk). Time stamp indicates minutes:seconds. (**G-H’**) R-GECO fluorescent intensity graphs are shown for single cells in contact with a cytoneme from GFP-expressing (**G** and **H**) or SHH-GFP-expressing (**G’** and **H’**) cells. Flux activity occurring over 15-minute contact periods in the absence (**G-G’**) or presence (**H-H’**) of the inverse SMO agonist cyclopamine is shown. (**I**) Flux rates per minute were calculated for 14-51 individual cells per condition. (**J**) Total flux time for R-GECO reporter cells is shown (n=27 for GFP and 15 for SHH contact). (**K**) Box plot (min/max whiskers) comparison of individual R-GECO cells (n=15) receiving SHH-GFP cytoneme deposits measured as proportion of time spent in flux in the absence of a SHH deposit, or within 20 seconds following deposit. All data are presented as mean ± SD, unless stated otherwise. ns = not significant, *** *p*<0.001.

The atypical actin-based motor Myosin 10 (MYO10) is thought to promote filopodial outgrowth through facilitating anterograde transport of protein cargo that supports growth and maintenance of the cellular structures (*9*–*13*). We tested for localization of MYO10 to the projections by expressing GFP tagged MYO10 in NIH3T3 cells. Indeed, MYO10-GFP was enriched in tips of projections from SHH-expressing cells, and showed co-localization with SHH puncta in a subset of the extensions (Figure 1C,C’). Live imaging revealed that the MYO10-tip-enriched extensions were highly dynamic and capable of forming stable connections with MYO10-positive extensions from neighboring cells (Movies 1-3). Shorter extensions that maintained contact with the growth substrate were largely immobile, suggesting they were likely adhesion or retraction fibers, rather than dynamic cytonemes (Movies 1, 2) (*7*, *10*).

Cytoneme-like structures were detected in the absence of SHH expression in both MEFs and NIH3T3 cells, indicating that the morphogen is not required for their initiation. However, SHH expression raised the cytoneme occurrence rate in both cell types from ~30% to ~60% (Figure 1D), and more than doubled the average number of cytonemes per NIH3T3 cell (Figure 1E). These results suggest that morphogen expression enhances cytoneme production or stability in murine cells, which is consistent with the established ability of HH to promote the stability of *Drosophila* cytonemes (*1*).

### Cytonemes deliver a SHH activation signal

To determine whether SHH-containing cytonemes transmitted an activation signal to receiving cells, we developed a contact-mediated activation assay in which SHH pathway induction in receiving cells could be rapidly detected. The current temporal indicator of pathway induction tracks accumulation of the SHH signal transducing G protein-coupled receptor Smoothened (SMO) into primary cilia (*14*). However, both active and inactive SMO proteins cycle through the primary cilium, making this assay sub-optimal for tracking SMO activation in real time (*15*, *16*). Active SMO signals through Gαi heterotrimeric G proteins to raise intracellular Ca^2+^, which we reasoned would be a rapid and activation-specific read-out for pathway induction resulting from cytoneme-based ligand delivery (*17*–*19*). To monitor Ca^2+^ flux, R-GECO-expressing Ca^2+^ reporter cells were co-cultured with NIH3T3 cells expressing SHH-GFP or GFP control. R-GECO cells in contact with cytonemes from GFP-positive cells were monitored for Ca^2+^ reporter flux (Figure 1F-G’, S2 and Movies 4-5). To increase the likelihood that signals would result from deposition of SHH via cytonemes, and not from SHH secreted into the media, media wash-out was performed at 15-minute intervals for the duration of data acquisition. We counted any R-GECO Ca^2+^ flux with a minimum peak fluorescence of 50 to be a positive event. This flux value was determined by examining the distribution of fluorescence intensity of R-GECO cells in contact with GFP control cells (n=27). A value of 50 was determined to be within the 90^th^-95^th^ percentile of flux value distributions in control cells, indicating any flux over 50 would likely be significantly above the control intensity range. Consistent with a SHH-induced signal, Ca^2+^ flux rates of R-GECO reporter cells in contact with SHH containing cytonemes were more than twice the rate of flux observed in reporter cells contacted by cytonemes from GFP-expressing control cells (Figure 1I). Furthermore, R-GECO cells in continuous contact with SHH-containing cytonemes had a significantly higher total time spent in flux than cells in contact with control cell cytonemes (Figure 1J). R-GECO reporter cells typically produced transient Ca^2+^ pulses within ~10-20 seconds of SHH-GFP release from cytonemes docked to their cell membranes (Figure 1G’, red dashes). To determine whether there was a statistically significant correlation between cytoneme-mediated ligand delivery and Ca^2+^ response, we documented all flux events with a mean intensity of over 50 that occurred within 20 seconds of SHH deposition (n=15 cells). A Wilcoxon signed rank test performed against these results confirmed that reporter cells spent a significantly greater amount of time in positive flux within a 20 second window following a SHH deposit than they did outside this response window (*p*=6.1e-05, Figure 1K). As such, a significant correlation between SHH delivery and Ca^2+^ release was confirmed. Importantly, SHH-stimulated Ca^2+^ flux was blocked by treatment with the inverse SMO agonist cyclopamine, confirming specificity of the Ca^2+^ response to SHH pathway activation (Figure1 H-I). Thus, cytonemes can deliver SHH to induce a *bona fide* SMO activation signal.

### SHH is trafficked inside cytonemes

Our observation that cytonemes appeared to transport distinct puncta of SHH toward target cells (Figure 1F) prompted us to investigate the mode by which morphogen reached the cytoneme tip. HHs are covalently modified by cholesterol at their carboxyl-termini, raising the possibility that they could travel along cytoneme membranes inside cholesterol-rich lipid rafts (*20*–*23*). To test for SHH localization to rafts along NIH3T3 cytoneme membranes, we assayed for ligand colocalization with a fluorescently-labeled cholera toxin (CTX) raft marker. Although rafts were evident along the length of cytonemes, they rarely contained SHH, as evidenced by a negative correlation coefficient between CTX and SHH signals (Figure S4A-B). Thus, SHH is unlikely to transport along cytoneme membranes in raft-like domains.

We next considered that SHH might load into vesicular structures for transport inside cytonemes because studies in both *Drosophila* and mouse suggest that HH ligands are released from producing cells in exosomes (*2*, *24*). To investigate this, we tested whether SHH localized inside cytonemes or to the outside leaflet of cytoneme membranes. Cells expressing SHH-GFP were subjected to extracellular immuno-staining with anti-SHH antibody prior to fixation. GFP fluorescence was used to track total SHH and antibody signal (ex-SHH) was used to monitor the ligand pool exposed to the extracellular environment. SHH-GFP and ex-SHH signals were both evident in puncta on the plasma membrane of SHH-expressing cells (Figure 2A, arrows). Notably, surface aggregates of ex-SHH evident along membranes of the cell body were rarely seen along cytonemes. Conversely, SHH-GFP signal was consistently detected in cytonemes, suggesting SHH is positioned inside cytoneme membranes (Figure 2A’, S3, S5A-B’). Consistent with this hypothesis, SHH co-localized with CD9-mCherry and CD81-mCherry exosomal markers along cytonemes (Figure 2B-C). Thus, ligand likely traffics inside cytonemes through a vesicular transport mechanism.

**Figure 2:**
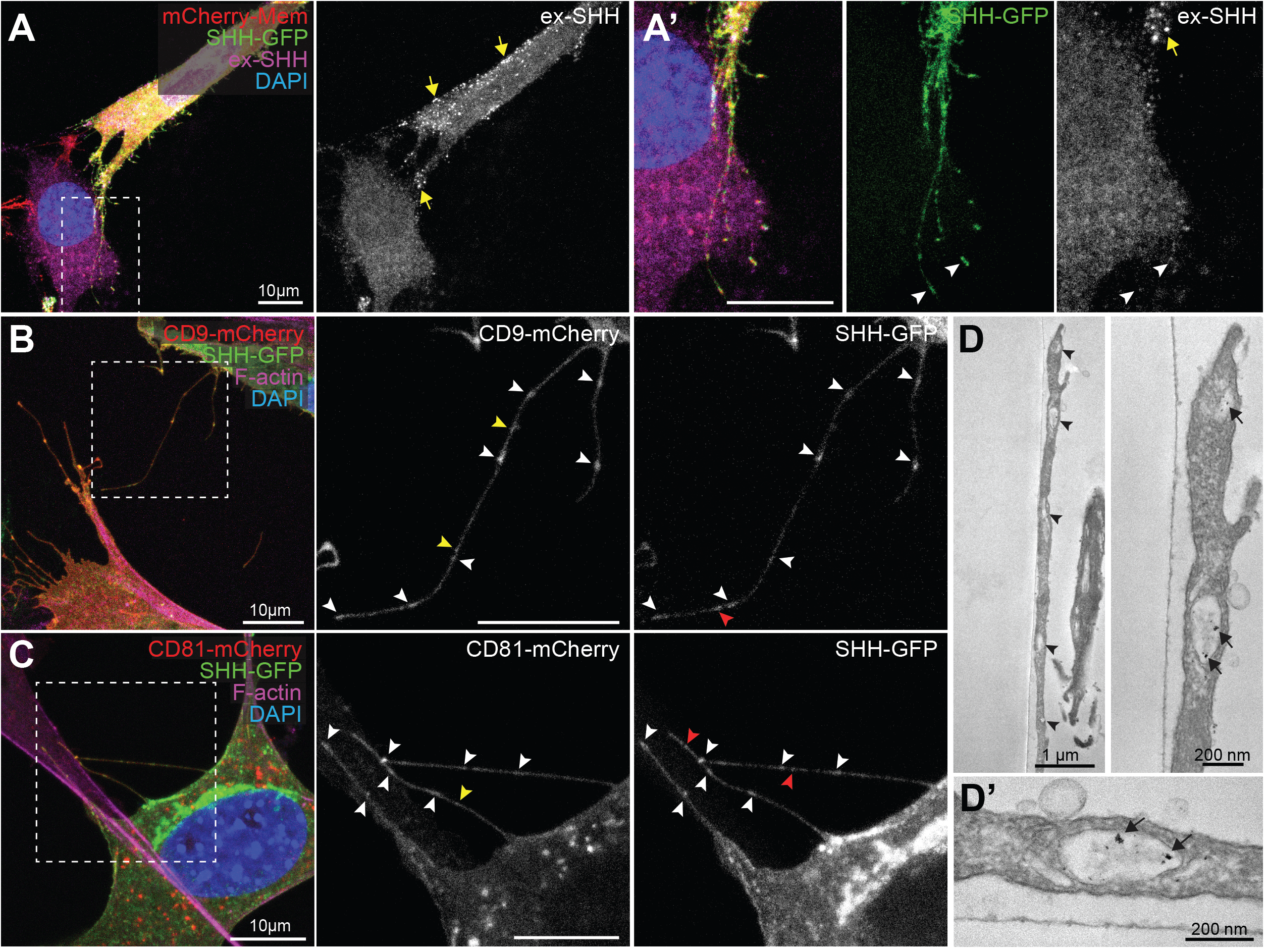
SHH is transported inside cytonemes. (**A,A’**) An NIH3T3 cell expressing SHH-GFP and mCherry-Mem is shown. Whereas SHH-GFP (green) is detected along cytonemes and at tips (**A’**, arrowheads), staining for extra-cellular SHH (ex-SHH, magenta and white) shows signal only on the cell body (**A-A’**, yellow arrows). (**B,C**) CD9-mCherry (**B**) and CD81-mCherry (**C**) colocalize with SHH-GFP in punctae along cytonemes (white arrowheads). CD9/CD81punctae lacking SHH are indicted by yellow arrowheads. SHH-GFP punctae lacking CD9/CD81 are indicted by red arrowheads. (**D,D’**) Transmission electron microscopy sections of a SHH-mCherry expressing cell cytoneme immunolabelled with anti-mCherry. Cytonemes contain vesicles (**D**, arrow heads), a subset of which contain SHH-mCherry (**D,D’** arrows). For all panels, nuclei are marked by DAPI (blue).

To evaluate whether SHH progressed through cytonemes in vesicles, SHH-mCherry-expressing NIH3T3 cells were examined by immuno-electron microscopy using anti-mCherry antibody. Transmission electron microscopy was performed on 70nm sections of cells and their cytonemes. Cytonemes of SHH-producing cells contained multiple vesicles (Figure 2D, arrowheads), many of which were positive for SHH-mCherry (Figure 2D,D’ arrows, Supplemental Figure 5). Although HH family ligands have been reported to enrich in RAB18 and CD63 containing exosomes in neuronal cells and *Drosophila* tissues, respectively, SHH failed to colocalize with these markers in cytonemes of NIH3T3 cells (Figure S4C-E)(*2*, *24*). Thus, cell-type specific vesicular loading may occur.

### Myosin 10 promotes cytoneme-based SHH transport

We hypothesized that if SHH undergoes vesicular trafficking inside cytonemes, a molecular motor would likely contribute to its movement. Because MYO10 co-localized with SHH at cytoneme tips (Figure 1C’), we tested whether inhibition of MYO10 would impact ligand movement along the specialized filopodia. MYO10-dependent effects on SHH cytoneme mobility were assayed by monitoring fluorescence recovery after photobleaching (FRAP) of the two proteins. Cytonemes of NIH3T3 cells expressing cytoplasmic GFP, mCherry-Mem, MYO10-GFP or SHH-mCherry were photobleached, and recovery of each protein to cytoneme tips was calculated in the absence or presence of ionomycin, which is proposed to attenuate MYO10 motor activity through raising intracellular Ca^2+^ (*25*, *26*). SHH-mCherry and MYO10-GFP cytoneme signals recovered at similar rates in vehicle-treated cells (~0.25±0.12 and 0.27±0.15 µm/s respectively), but failed to recover following ionomycin treatment. Conversely, membrane diffusion rates, which were calculated by monitoring mCherry-Mem recovery to cytoneme tips, were not reduced by ionomycin treatment (Figures 3A-B, S6, Movies 6 and 7). Recovery of cytoplasmic GFP signal to cytoneme tips was so rapid we were unable to calculate accurate recovery rates in either condition. These results suggest SHH is unlikely to travel along cytonemes through cytoplasmic or membrane diffusion-based mechanisms, and is instead actively transported along the specialized filopodia, potentially by MYO10.

**Figure 3:**
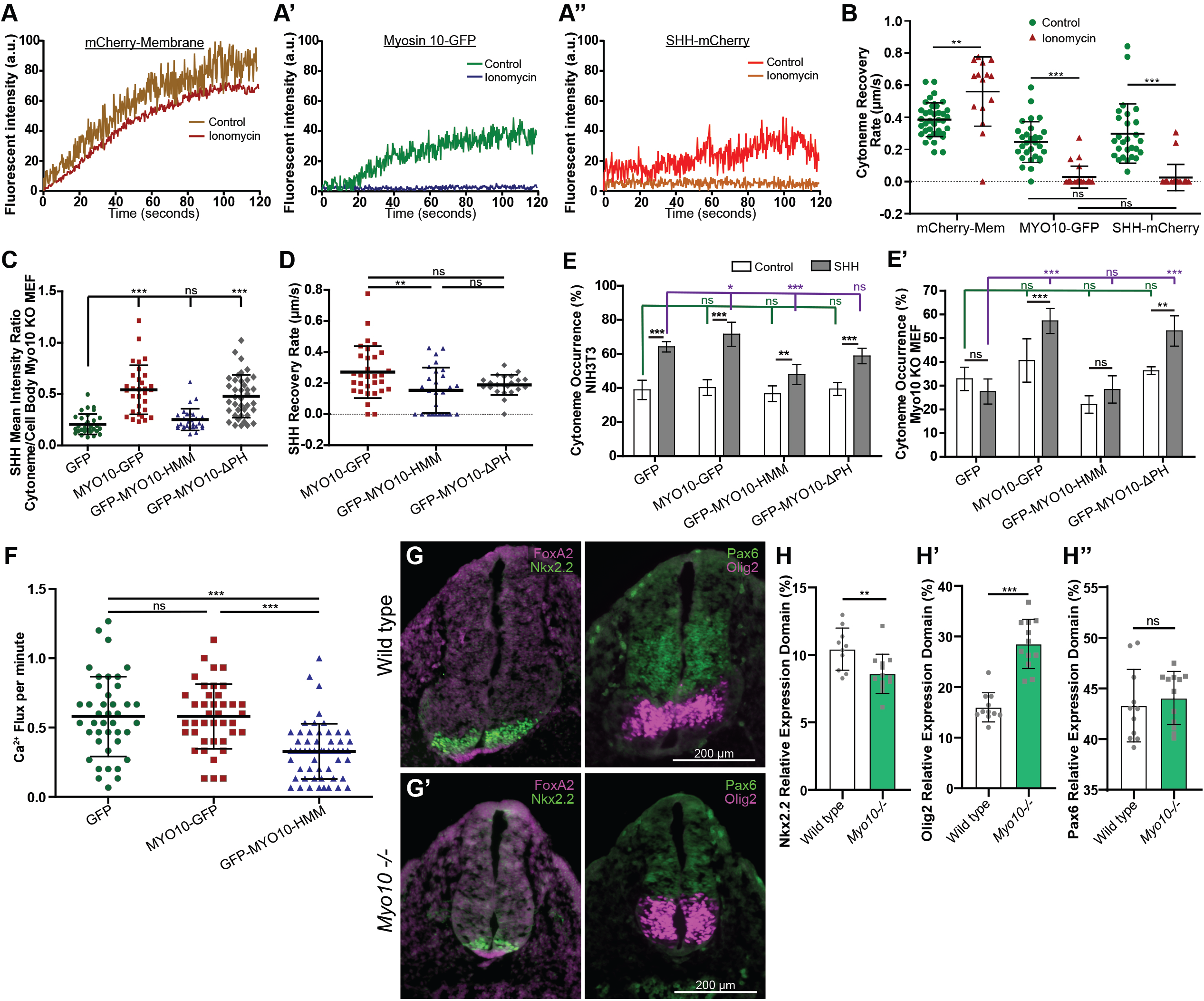
Myosin10 facilitates cytoneme-based SHH transport and Delivery. (**A-A”**) NIH3T3 cells expressing the indicated fluorescent proteins were subjected to FRAP. Representative FRAP curves of (**A**) mCherry-Mem, (**A’**) MYO10-GFP, and (**A”**) SHH-mCherry to a cytoneme tip in control (DMSO) or ionomycin-treated (2.5 µM) conditions are seen. (**B**) Scatter plots show the recovery rates of the indicated fluorescent proteins to cytoneme tips in control (DMSO) or ionomycin-treated conditions, calculated from FRAP curves (n=14-38 cells). (C) Scatter plots of mean cytoneme to cell body SHH fluorescent signal ratios in *Myo10-/-* MEFs co-expressing GFP or the indicated MYO10 proteins (n=28-38). (**D**) Scatter plots of FRAP calculated recovery rates for SHH-mCherry movement toward cytoneme tips in NIH3T3 cells co-expressing the indicated MYO10 proteins (n=22-32). (**E,E’**) Cytoneme occurrence rates were calculated for MEM-fixed NIH3T3 and *Myo10-/-* MEFs co-expressing mCherry-Mem (control) or SHH plus the indicated MYO10 proteins or GFP control. (**F**) Ca^2+^ flux rates per minute were determined for R-GECO reporter cells in contact with cytonemes of SHH producing cells co-expressing the indicated MYO10 proteins (n=40-55 cells per condition). (**G,G’**) Representative sections of posterior neural tube from E10.5 wild type and *Myo10-/-* mice were immuno-stained for FOXA2 (magenta, left), NKX2.2 (green, left), PAX6 (green, right), and OLIG2 (magenta, right). (**H-H”**) Expression domains of (**H**) *Nkx2.2*, (**H’**) *Olig2*, and (**H”**) *Pax6* in wild type or *Myo10-/-* embryos were calculated relative to the dorso/vertral length of the neural tube. Dots represent the individual sections examined from the representative embryo. Three embryos per condition were analyzed. All data are presented as mean ± SD. ns = not significant, * *p*<0.05, ** *p*<0.01, *** *p*<0.001.

To test whether SHH enrichment in cytonemes required MYO10 function, we generated MYO10-null MEFs from *Myo10^m1j/m1j^* mutant mice, and assessed SHH cytoneme dynamics in this genetic background (*27*). SHH was expressed in MYO10 mutant MEFs, and cytoneme to cell body SHH signal intensity ratios were determined (Figures 3C). MYO10 mutant MEFs exhibited low SHH cytoneme to cell body signal intensity ratios, indicating inefficient cytoneme enrichment of the morphogen in the absence of MYO10. Enrichment of SHH in cytonemes was restored by co-expression of wild type or pleckstrin homology domain-deficient (ΔPH) MYO10, but not by a MYO10 mutant lacking its cargo binding domains (MYO10-HMM) (*28*). Comparable effects on SHH cytoneme enrichment were seen in wild type NIH3T3 cells over-expressing these MYO10-GFP variants. Whereas wild type and MYO10ΔPH did not alter the ratio of SHH or SHH-mCherry in cytonemes, MYO10-HMM over-expression reduced SHH cytoneme localization, and attenuated SHH-mCherry FRAP to cytoneme tips (Figure 3D, S7A-F). All MYO10 variants showed similar cytoneme FRAP rates, suggesting that failure of SHH to be transported by MYO10-HMM was likely due to compromised cargo binding, and not due to defective MYO10-HMM motor activity (Figure S7G).

MYO10 contributes to the formation and maintenance of filopodia, and SHH increases cytoneme occurrence rates in NIH3T3 cells and wild type MEFs (Figure 1D,E) (*9*). Hence, we tested for a role for MYO10 in SHH-stimulated cytoneme occurrence in wild type and *Myo10* mutant cells. In NIH3T3 cells, expression of wild type MYO10 modestly enhanced the ability of SHH to increase cytoneme occurrence rates over baseline. MYO10-HMM suppressed occurrence rates (Figure 3E), which we speculate resulted from the mutant protein oligomerizing with endogenous MYO10 to disrupt its ability to bind and transport cargo (Figure 3E and S6G). In *Myo10-/-* MEFs, SHH failed to stimulate cytoneme occurrence. Cytoneme occurrence rate increases in the presence of SHH were rescued by reintroduction of either wild type MYO10 or MYO10-ΔPH, but not by MYO10-HMM (Figure 3E’), further supporting an essential role for the cargo domain for SHH cytoneme biology. Consistent with this hypothesis, co-expression of GFP-MYO10-HMM with SHH in ligand-producing NIH3T3 cells reduced SHH-induced Ca^2+^ flux in co-cultured R-GECO reporter cells to significantly lower rates than those observed upon co-culture with ligand-producing cells co-expressing GFP or wild type MYO10-GFP with SHH (Figure 3F). Combined, these results suggest MYO10 is required for the cytoneme-promoting effects of SHH, and also for cargo domain-mediated transport of SHH to cytoneme tips for delivery to target cells.

### Myosin 10 promotes SHH signaling in vivo

*Myo10 -/-* mice are semi-lethal with ~60% of homozygous mutants exhibiting exencephaly with embryonic or perinatal lethality. Surviving animals display white belly spots, with a subset of these animals also exhibiting syndactyly (*27*, *29*). Exencephaly and syndactyly can be attributed to reduced Bone Morphogenic Protein (BMP) signaling and de-repression, rather than disruption, of SHH signaling (*30*, *31*). These phenotypes are seemingly inconsistent with our *in vitro* observations that MYO10 promotes cytoneme stability and transport of SHH (Figure 3A-F). Therefore, in an effort to understand the effects of MYO10 loss on SHH signaling *in vivo*, we analyzed developing neural tubes from *Myo10^m1J/m1J^* E10.5 embryos (*27*). SHH is expressed in the notochord and floor plate of the developing neural tube, and signals in a ventral to dorsal trajectory to specify distinct neural progenitor domains. This occurs through SHH inducing expression of cross-repressive transcription factors *Nkx2.2* and *Olig2*, and repressing *Pax6*. Induction of these genes is exquisitely sensitive to alteration of SHH signaling, so monitoring their expression allows for robust analysis of gradient function (*32*, *33*). Examination of wild type and *Myo10^m1J/m1J^* neural tubes revealed normal floorplate specification in *Myo10* mutants, as evidenced by similar *FoxA2* expression domains between the two genotypes (Figure 3G, G’). However, the progenitor domain marked by the high-threshold SHH target gene *Nkx2.2* was compressed in *Myo10* mutants, allowing for ventral expansion of the progenitor domain marked by the intermediate SHH target *Olig2* (Figure 3G-H”). *Pax6* expression was not overtly altered in *Myo10* mutants, suggesting that loss of MYO10 function specifically compromised induction of high-threshold SHH-activated target genes (Figure 3G-H’’). Similar patterning alterations were evident in more anterior sections of *Myo10* mutant neural tubes, albeit to a lesser extent (Figure S8). The failure of MYO10 mutants to exhibit strong SHH loss-of-function phenotypes is not unprecedented given the variable phenotypic penetrance observed in *Myo10* knockout animals (*27*). We speculate that functional compensation by cytoneme-independent mechanisms of SHH distribution likely occur to limit the impact of MYO10 loss on the SHH morphogen gradient establishment (reviewed in Hall et al. (2019) (*34*)). Given the established role for cytonemes in transport of the *Drosophila* BMP ortholog, Decapentaplegic (DPP) (*35*), we further speculate that pronounced exencephaly observed in some murine *Myo10* mutants is likely due to compromised BMP signaling, and not the result of SHH de-repression following MYO10 loss.

### SHH co-localizes with DISP and coreceptors in cytonemes

Having identified MYO10 as a new functional player in cytoneme occurrence and SHH transport, we next wanted to determine whether known SHH-binding partners would also impact cytoneme occurrence in mammalian cells. Experiments in *Drosophila* and chick model systems suggest a role for the SHH deployment protein DISP and co-receptors BOI/BOC and iHOG/CDON in cytoneme function (*1*–*3*, *21*, *36*). In flies, DISP promotes cytoneme stability of ligand producing cells, and in chick, BOC stabilizes cytonemes of SHH receiving cells (*1*, *3*). Because iHOG/CDON has been reported to stabilize and localize to exovesicles in cytonemes of ligand-producing cells in flies (*2*, *36*), we hypothesized that DISP and CDON or BOC might function together to influence cytoneme occurrence or function in mouse cells. DISP-HA and GFP-tagged BOC or CDON were co-expressed in NIH3T3 cells in the absence and presence of SHH, and colocalization between the three proteins was assessed. Confocal microscopy revealed that all three proteins localized to cytonemes, but that DISP did not significantly co-localize with either BOC or CDON along cytoneme membranes in the absence of SHH (Figure 4A,D,F,G). Co-expression of SHH increased colocalization between BOC and DISP throughout cytoneme membrane and in SHH-positive puncta (Figure 4A-B’, F). Notably, puncta containing all three proteins were evident in cells abutting SHH-containing cytonemes (Figure 4B and zoom in B’, arrowheads). As such, DISP and BOC may be released to target cells along with ligand, as has been reported for iHOG in *Drosophila* (*2*). Consistent with ligand-containing endosomes being internalized by receiving cells, immunoelectron microscopy revealed early and late-endosomal structures containing SHH near cytoneme contact points on the signal-receiving cell (Figure 4C).

**Figure 4:**
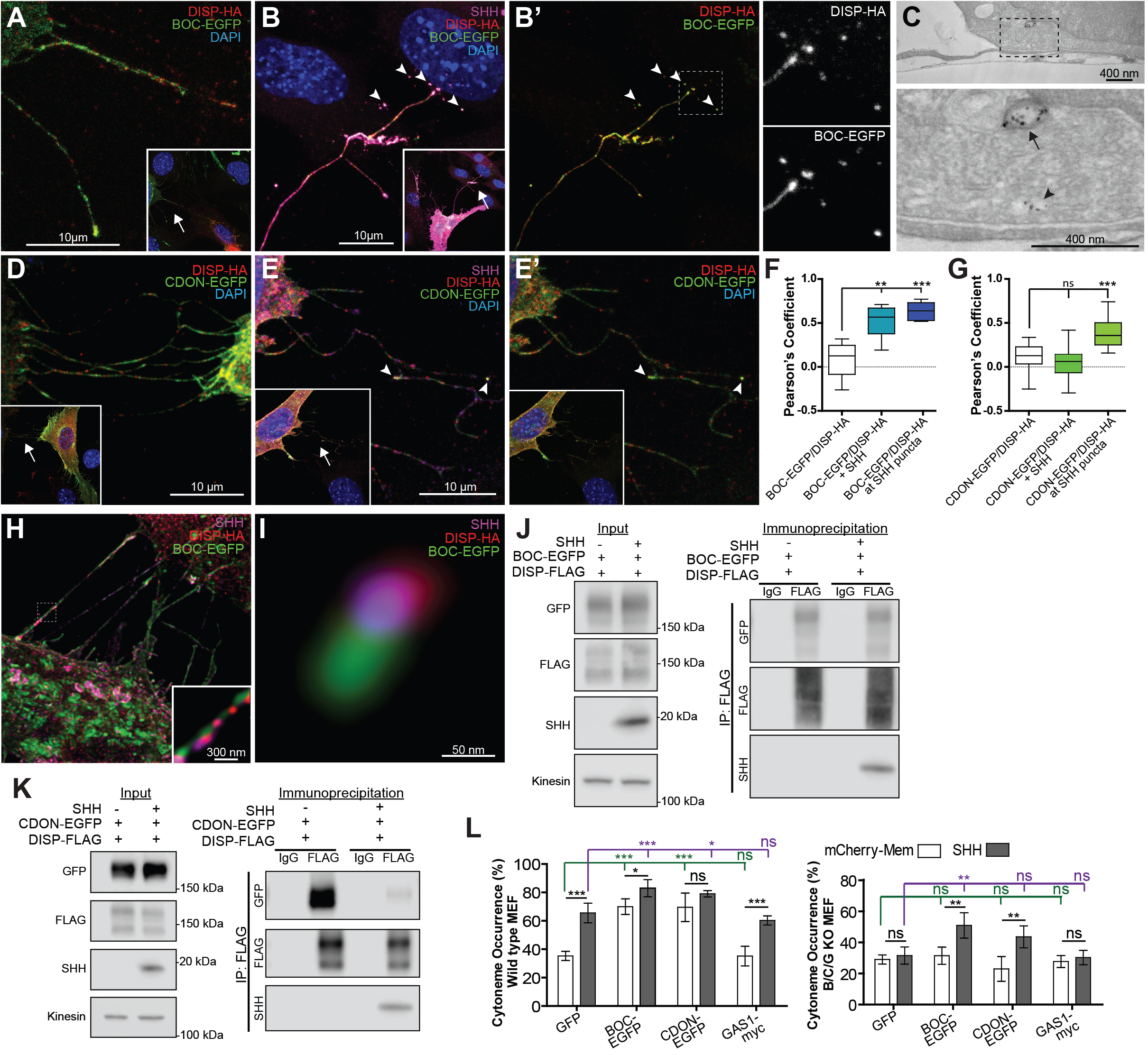
DISP, BOC and CDON ligand complexes influence SHH cytonemes. (**A**) DISP-HA (red) and BOC-EGFP (green) localize to cytonemes of NIH3T3 cells. Inset shows lower magnification, with arrow indicating magnified area. (**B-B’**) SHH (magenta)-expressing NIH3T3 cell cytonemes show co-localization between DISP-HA and BOC-EGFP. Inset shows lower magnification, with arrow indicating magnified area. Punctae visible on signal-receiving cells contain BOC-EGFP and DISP-HA (arrowheads and right). Dashed lines indicate magnified regions, right. (**C**) TEM section of a SHH-mCherry expressing cell cytoneme in contact with a receiving cell. SHH is immunolabeled with anti-mCherry and is present in early (arrow head), and late (arrow) endosomal compartments of the receiving cell near the cytoneme contact point. (D) DISP-HA and CDON-EGFP (green) are present in cytonemes of NIH3T3 cells. Inset shows lower magnification, with arrow indicating magnified area. (**E-E’**) DISP-HA, CDON-EGFP, and SHH localize to cytonemes of NIH3T3 cells. Arrow heads identify colocalization. (**F-G**) Box plots of Pearson’s correlation coefficient value (min/max whiskers) measuring colocalization between (**F**) DISP-HA and BOC-EGFP, or (**G**) DISP-HA and CDON-EGFP under the indicated conditions. (n= >30 cytonemes per condition). (**H-I**) Stimulated emission depletion microscopy images of SHH, BOC-EGFP, and DISP-HA in cytonemes (**H**) and a representative magnified puncta (**I**). (**J-K**) DISP-FLAG was immunoprecipitated from lysate from NIH3T3 cells expressing DISP-FLAG and BOC-EGFP (**J**) or CDON-EGFP (**K**) in the absence or presence of SHH. Lysate input is shown at left and FLAG immunoprecipitates are shown at right. IgG serves as control. (**L**) Cytoneme occurrence rates for wild type and *Boc/Cdon/Gas1-/-* MEFs expressing mCherry-Mem or SHH in the presence of GFP or the indicated SHH co-receptor. All data are presented as mean ± SD, unless stated otherwise. ns = not significant, * *p*<0.05, ** *p*<0.01, *** *p*<0.001.

To better understand the associations between cytoneme-localized DISP, BOC and SHH, stimulated emission depletion (STED) microscopy was used to examine individual SHH puncta within cytonemes. STED showed DISP and SHH consistently positioned adjacent to BOC, suggesting that a trimeric complex may occur in cytonemes in the presence of ligand (Figure 4H, I). To test for interaction between the three proteins, co-immunoprecipitation experiments were performed using lysates from NIH3T3 cells expressing DISP-FLAG and BOC-EGFP in the absence and presence of SHH. BOC-EGFP was captured on anti-FLAG beads in the absence of SHH, suggesting that DISP and BOC can associate (Figure 4J, and S9A). Upon ligand expression, SHH incorporated into DISP-FLAG/BOC-EGFP immunocomplexes without significantly altering BOC binding. Thus, SHH is not required for biochemical association between DISP and BOC, but may promote enrichment of the trimeric complex in cytonemes (4A-B’, J).

BOC and CDON are semi redundant for co-receptor function in PTCH-SHH binding, but do show differential expression and functionality in temporal and tissue specific contexts (*37*–*41*). Likely consistent with context-dependent functionality, co-localization dynamics between DISP and CDON differed from what was observed for DISP and BOC. CDON colocalized with DISP at SHH-positive puncta in cytonemes, albeit to a slightly lesser extent than did BOC (Figure 4D-G). However, unlike what was observed for BOC, ligand expression did not increase co-localization between DISP and CDON along the length of cytonemes (Figure 4F, G). Biochemical interrogation of DISP-CDON binding by immunoprecipitation analysis revealed that whereas CDON-GFP was captured on FLAG beads by DISP-FLAG in the absence of ligand, its association with DISP-FLAG was reduced upon SHH-DISP binding (Figure 4K, S9B). Thus, distinct functional pools of CDON with differential affinity toward DISP may exist.

### BOC and CDON promote SHH cytoneme formation

*Drosophila* iHOG (CDON) and chick BOC proteins have been reported to localize to cytonemes and influence their behavior *in vivo* (*3*, *21*, *36*). To determine the effects of BOC and CDON on cytonemes of SHH-expressing murine cells, they were co-expressed with SHH in MEFs, and cytoneme occurrence rates were determined. The vertebrate-specific SHH co-receptor GAS1, which has not yet been investigated for a role in cytonemes, was also tested (*42*). GAS1 expressing cells showed baseline and SHH-induced cytoneme occurrence rates similar to GFP-expressing control cells, indicating that GAS1 over-expression does not actively promote or stabilize cytonemes in the absence or presence of ligand (Figure 4L). Conversely, both BOC and CDON expression elevated baseline cytoneme occurrence rates near to SHH-stimulated levels, and modestly enhanced the ability of SHH to increase occurrence (Figure 4L). To determine whether BOC, CDON, GAS1 or a combination of the co-receptors was required for SHH-induced cytoneme occurrence rate increases, we expressed SHH in *Boc-/-, Cdon-/-, Gas1-/-* triple KO MEFs (B/C/G KO) (*37*), and quantified occurrence rates in control and co-receptor re-expressed conditions (Figure 4L, right panel). GFP-expressing B/C/G KO cells failed to increase cytoneme occurrence upon SHH expression, indicating that at least one of the co-receptors is required to facilitate SHH-induced cytoneme occurrence. GAS1 re-expression did not rescue the ability of SHH to promote cytoneme occurrence, further supporting that GAS1 is not a potent modulator of SHH cytoneme function. Conversely, re-expression of either BOC or CDON rescued SHH-mediated cytoneme occurrence increases in the triple KO cells (Figure 4L). Thus, we conclude that either BOC or CDON is required for SHH-induced cytoneme biogenesis or stability, and that this functionality is likely conferred by co-receptor interactions with DISP/SHH complexes in the specialized filopodia. We do not know the mechanism(s) by which SHH-containing co-receptor complexes promote cytoneme induction or stability. However, the reported ability of BOC to activate the cytoskeletal regulator JNK during neuronal differentiation (*43*), suggests that BOC or CDON might connect SHH with actin remodelers for cytoneme movement.

By analyzing cytonemes of cultured murine cells, we were able to interrogate the molecular mechanisms contributing to cytoneme-based morphogen transport. Our investigation revealed, for the first time, that release of SHH from cytoneme tips induces a rapid, SMO-dependent signal response in target cells. Our results suggest that SHH may also signal in an autocrine manner in ligand-producing cells to promote cytoneme biogenesis or stability through a complex containing the deployment protein DISP and co-receptor BOC or CDON. SHH cytoneme stability and morphogen delivery are also promoted by the molecular motor MYO10, which facilitates vesicular trafficking of SHH inside the dynamic structures. Importantly, loss of MYO10 altered SHH-dependent tissue patterning *in vivo*, supporting that analysis of cultured cell cytonemes can predict biologically relevant contributors to morphogen gradient function during development. The proven utility of cultured cells for analyzing cytoneme biology reveals that *in vitro* systems can function as tractable models for interrogating morphogen transport. Furthermore, cultured cells may also allow for investigation of how cytonemes synergize with other morphogen dispersion processes to ensure gradient robustness during tissue development.

## Methods and Materials

### Immunofluorescence and imaging

Cell fixation and staining were performed using MEM-fixation protocols (*6*). The following antibodies and dilutions were used: rabbit anti-SHH (H-160) (1:100; Santa Cruz), mouse anti-GFP (4B10) (1:500; CST), rat anti-HA (1:250; Roche), mouse anti-CD63 (E-12) (1:100; Santa Cruz), rabbit anti-Myc-Tag (2272) (1:400; CST). Secondary antibodies (Jackson ImmunoResearch and Invitrogen) were used at a 1:1000 dilution. Lipid raft staining was performed using Cholera Toxin Subunit B (Recombinant) (CTX), Alexa Fluor 488 Conjugate (Invitrogen). CTX was dissolved in chilled PBS to a final concentration of 1.0 mg/mL. CTX was incubated with cells for 20 minutes at 4°C to prevent endocytosis. Cells were rinsed 3 times in chilled PBS prior to MEM-fixation. Extracellular staining was performed by diluting antibodies in 4°C PBS supplemented with 5% normal goat serum. Antibody solutions were then incubated for 30 minutes on live cells on ice to prevent endocytosis. Cells were rinsed 3 times in chilled PBS prior to MEM-fixation. Microscopy images were taken with a TCS SP8 STED 3X confocal microscope (Leica) for fixed and live cell imaging.

### Fluorescence Recovery After Photobleaching (FRAP)

FRAP assays were carried out on a Bruker Opterra swept field confocal microscope, equipped with an enclosure box at 5% CO_2_ and imaging and objective heater at 37°C. NIH3T3 cells were imaged in phenol red-free standard growth media. For assays involving ionomycin, standard growth media was replaced immediately prior to imaging with phenol red- and serum-free media supplemented with 0.083% DMSO (control), or DMSO with 2.5 µM ionomycin (#9995, CST). Image acquisition was performed with 60x/1.4NA/Oil objective lens (CFI Plan Apo Lambda) with a 30 µm pinhole array and 70 µm width slit. Fluorescence was recorded with a 5 second (s) baseline followed by a complete photobleaching of a cytoneme with 488-nm and 561-nm lasers. Fluorescence recovery was recorded for 120 s with 100 ms exposure per channel with frames taken every 277 ms. Fluorescence recovery of individual regions of interest (ROI) along the cytoneme to its tip were normalized with pre-FRAP equal to 100% and post-FRAP equal to 0%. FRAP curves were corrected for any loss of fluorescence during acquisition (*44*).

### Generation of cell lines and culture

Cells were cultured at 37°C in 5% CO_2_. NIH3T3 (CRL-1658), HEK293T (CRL-11268) and LightII (JHU-68) cells were obtained from ATCC. *Boc/Cdon/Gas1 -/-* MEFs were obtained from B. Allen (*37*). *Myo10-/-* MEFs were generated from mice obtained from and cryo-recovered by The Jackson Laboratory (stock number 024583, B6.Cg-Myo10m1J/GrsrJ). MEFs were generated as previously described (*45*). Briefly, pregnant dams were harvested at E12.5-13.5 and embryos were dissected in 1X PBS, then decapitated and internal organs removed. The remaining tissue was rinsed in 1X PBS, then finely minced into pieces in a dish containing Trypsin-EDTA (0.25%) (Gibco). The dish was placed at 37°C in an incubator for 15 minutes, then an additional 2 mL of Trypsin-EDTA was added, tissue was vigorously pipetted, then placed back in the incubator at 37°C for an additional 10 minutes. The solution was transferred to a 15 mL conical tube and contents were allowed to settle for 2 minutes. Supernatant was removed, and then centrifuged for 5 minutes at 200 × g. The cell pellet was resuspended in MEF media (see below) and plated in a 60 mm plate and left overnight. Each line was then SV40-transformed and single cell selection was performed by serial dilution in a 96 well plate. MEF lines were derived from 5 different *Myo10* mutant embryos and 5 wild type littermates.

Cells were maintained in DMEM (Life Technologies) supplemented with 10% bovine calf serum (Fisher Scientific) and 1% Penicillin Streptomycin solution (Gibco). HEK293T cells utilized 10% heat-inactivated fetal bovine serum (Corning). Cell lines were routinely validated by functional assay and western blot as appropriate and tested monthly for mycoplasma contamination by MycoAlert (Lonza). Transfection of plasmid DNA was performed with Lipofectamine 3000 and P3000 reagent (ThermoFisher Scientific), according to manufacturer’s instructions. When required, the final amount of DNA used for transfection was kept constant by the addition of control vector DNA. All cells were harvested 36 h after transient DNA plasmid transfection for subsequent assays.

### In vivo analysis

Wild type and *Myo10^m1J/m1J^* embryos were harvested and processed for immunohistochemistry between stages E9.5 to E12.5. Pregnant dams were harvested, uterine horns removed, and embryos were dissected in 1X PBS, then rinsed three times. Embryos were fixed overnight at 4C in 2% PFA. The following day, embryos were rinsed three times in 1X PBS and moved to 30% sucrose to cryo-protect. The following day, embryos were frozen in O.C.T. Compound (Tissue-Tek) on dry ice. Embryos were sectioned transverse at 10 µm thickness on a Leica Microm CM1950 cryo-stat. Sections were briefly dried, then washed in 1X TBST, then blocked with 2% BSA, 1% goat serum, 0.1% Triton-X-100 in 1X PBS. Antibodies were diluted in blocking buffer and incubated overnight on sections at room temperature. The following antibodies and dilutions were used: mouse anti-FOXA2 (1:50; DSHB), rabbit anti-NKX2.2 (1:60; Novus), mouse anti-PAX6 (1:25; DSHB), and rabbit anti-OLIG2 (1:300; Millipore). Primary antibody was removed, sections were washed with 1X TBST three times, then incubated for 3 hours in secondary antibodies (Invitrogen) used at a 1:500 dilution. Sections were washed 3 times in 1X TBST, then rinsed with tap water, and cover slips were applied with ProLong Diamond mounting media. Sections were imaged on a Leica DMi8 widefield microscope and processed using LAS X. Relative expression domains were calculated by the mean length (dorso/ventral axis) per section of the target protein across the entire neural tube length (n>10 sections per condition).

### Ca^2+^ Flux assay

Approximately 0.4×10^6^ NIH3T3 cells were seeded into individual wells of 6-well plates one day prior to transfection. A total of 2µg plasmid DNA was transfected into individual wells. pCMV-R-GECO1 for ‘receiving’ sensor cells, and the appropriate construct combination (e.g. SHH-mCherry, GFP, MYO10-GFP, etc…) for the ‘producing’ cells. Six hours after transfection, cells were trypsinized and replated into 8 well, polystyrene chambers on a 1.5 borosilicate coverglass, 0.7 cm^2^/well (Nunc Lab-Tek II). Prior to cell addition, chamber wells were divided in half with #0 (0.08-0.13 mm) thick coverslips (Electron Microscopy Sciences) cut to size, with vacuum grease (Dow Corning) added along the lateral edges to retain a liquid-tight barrier. Receiving and producing cells were seeded onto opposite sides of the barrier and allowed to recover overnight. The following day the barrier was removed 3 hours prior to imaging allowing sufficient time for cells to migrate and cytonemes extend between producing and receiving cells. Media was removed prior to imaging, and cells were gently washed in PBS. Immediately prior to any imaging event, media was replaced to remove secreted SHH. In experiments where SMO activation was inhibited, 10 µM cyclopamine (LC Laboratories) was added to R-GECO1 cells 16 h prior to imaging.

Live imaging was performed at 37°C, 5% CO_2_ with resonant scanning for 15 minutes per area over the entire cytoneme/s depth (~4-6 µm), with Z-steps of ~0.6-1.0 µm. Maximum intensity projections were generated for subsequent analysis of the time-lapses.

### SHH deposits and Ca^2+^ flux quantification

Time-lapses of cells were analyzed using LAS X (Leica). SHH-GFP deposits onto R-GECO sensor cells were recognized if a SHH puncta was detected traversing a cytoneme from a producing cell body to accumulate at the tip, where in a successive frame fluorescence was absent and did not undergo retrograde movement. Puncta were identified by a fluorescent intensity signal >50% than background cytoneme fluorescence by single line scan along a cytoneme. R-GECO fluorescent intensity histograms of individual cells were normalized for each cell with minimum fluorescence equal to 0 and maximum to 100. Ca^2+^ flux occurrence was quantified as a relative peak in R-GECO fluorescence within a ~20 second window with a minimum peak fluorescence of 50. Maintained fluorescence over 20 seconds was considered a single flux. Total flux time was determined using a threshold to define an increased flux (i.e. a peak) using the R-GECO cells in contact with GFP-control samples. The threshold was determined to be a flux value of greater than 50 which lies between the overall 90th and 95th percentiles of the control samples. Next, the proportion of time (seconds) when flux values were greater than 50 was calculated for each control and SHH case sample. The proportion of time when flux values were greater than 50 was compared between cases and controls using the Wilcoxon rank sum test. For SHH cases only, the proportion of time when flux values were greater than 50 was compared by the occurrence of a SHH deposit (yes vs. no) using the Wilcoxon signed rank test. Twenty seconds was considered a biologically relevant time frame in which a deposit and a subsequent increased flux should occur (*18*, *46*). Statistical analyses were conducted using SAS software version 9.4 (SAS Institute, Cary, NC) and R version 3.6.0 (R Foundation for Statistical Computing, Vienna, Austria). A two-sided significance level of p<0.05 was considered statistically significant. R-GECO cells in contact with cytonemes that did not exhibit a single flux during the time-lapse were excluded from analysis.

### Image processing, measurements, and statistical analysis

#### Image processing

Following image acquisition, images were processed using LAS X (Leica), and Photoshop 2019 (Adobe), and figures were made using Illustrator 2019 (Adobe). Stimulated emission depletion images underwent deconvolution using automated sampling with Huygens Professional software (Scientific Volume Imaging). Videos were processed with ImageJ and Imaris (Bitplane). Images are presented as maximum intensity projections (MIP) of the z-stack acquisition spanning the cell, unless stated otherwise.

#### Cytoneme metrics

For quantification we defined cytonemes in cultured cells as cellular projections approximately < 200 nm in diameter, with a minimum length of 10 µm. Any cellular protrusions that originated from the basal surface of the cell and maintained continuous contact with the coverslip were excluded from analysis (*6*). For live imaging cytonemes were identified as motile protrusions, capable of elongation with the exception if a cytoneme was in contact with a nearby cell body or other cells’ cytonemes. For occurrence rate counts, a minimum of 100 cells per condition were counted, performed on triplicate coverslips with a minimum of two biological replicates.

#### Calculation of Diffusion/Recovery rates of proteins to cytoneme tips

Protein recovery rates to the cytoneme tip were derived by average velocity,

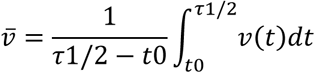

based upon the following conditions. 1) A cytoneme diameter is ~100 nm, below the diffraction limited resolution of the confocal microscope in which the FRAP data was acquired. Therefore, all data may be reduced to a single plane (a 1-dimensional line). 2) Photobleaching of the entire cytoneme allows for a single vector recovery from the cell body, as such 2D diffusion coefficients calculations are not required. FRAP data was analyzed by Igor Pro 8 (WaveMetrics) to calculate halftime recovery (τ_1/2_) of 2-3 ROIs along an individual cytoneme. τ_1/2_ is dependent upon the distance from the cell body, allowing for instantaneous velocity calculation.

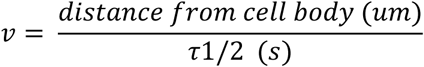

Values were then averaged per cytoneme. 16-39 individual cytonemes were analyzed for each condition.

#### Colocalization analysis

Image analysis was performed using CellProfiler (*47*). An image analysis pipeline was constructed to mask and isolate cytonemes. Output images were then run through a secondary pipeline measuring Pearson’s correlation coefficient between pixels of different fluorophores to calculate the relative colocalization of the proteins of interest within a cytoneme. A third pipeline was used for determining protein colocalization of 2 proteins at SHH puncta in cytonemes. This pipeline used the SHH pixels as a reference point within cytonemes to measure Pearson’s correlation coefficient between pixels of the other fluorophores of interest in contact with SHH. A minimum of 30 cytonemes were analyzed per condition.

#### Statistical analyses

All analyses were performed using GraphPad Prism. One-way analysis of variance was performed for multiple comparisons, with Tukey’s multiple comparison as a posttest. Significant differences between two conditions were determined by two-tailed Student’s t tests. All quantified data are presented as mean ± SD, with p < 0.05 considered statistically significant. Significance depicted as *p < 0.05, **p < 0.01, ***p < 0.001, ns = not significant.

#### Plasmid constructs

The following plasmids were used in this study: pCDNA (control vector) (Clontech), pCDNA3-EGFP (Addgene Plasmid #13031), pCMV-mCherry-Mem (Addgene Plasmid #55779), pCMV-R-GECO1 (Addgene Plasmid #32444), pCMV-mCherry-CD9 (Addgene Plasmid #55013), pCMV-mCherry-CD81 (Addgene Plasmid #55012), pEGFP-CD63-C2 (Addgene Plasmid #62964), pCMV-EGFP-Rab18 (Addgene Plasmid #49550), pEGFP-C1-hMyoX (Addgene Plasmid #47608), pCMV-mCherry2, pCDNA3-mSHH-FL, pCDNA3-mSHH-N, pCDNA3-mSHH-FL-EGFP, pCDNA3-mSHH-FL-mCherry2, pCDNA3-V5-Disp-HA, pCS2-hBOC-EGFP and pCS2-hCDON-EGFP (a gift from A. Salic), pEGFP-C2-bMyo10-HMM (*28*) and pEGFP-C2-bMyo10-Δ3PH, a modified version of pEGFP-bMyo10 where the 3 PH domains were removed via deletion of aa 1168-1491.

For the generation of fluorescently tagged SHH, GFP or mCherry2 was introduced into the SHH protein immediately 3’ to amino acid Gly198 with the addition of 10 amino acids (Alanine188 - AENSVAAKSG - Glycine197) downstream of GFP or mCherry2 including the intein cleavage-cholesterol attachment site, similar to what was previously descried Chamberlain et al., (2008)(*48*). Briefly, a Bgl2 site was introduced into SHH-FL after Gly198 by Quikchange (Agilent) using primers (forward 5’ GTGGCGGCCAAATCCGGCGGCAGATCTGGCTGTTTCCCGGGATCCGCC and reverse 5’ ggcggatcccgggaaacagccagatctgccgccggatttggccgccac). The 10 amino acid duplication 3’ to GFP or mCherry2 on SHH protein was introduced using Infusion (Clontech) using primers (forward 5’TCCGGCGGCAGATCTGCAGAGAACTCCGTGGCGGCCAAATCCGGCGGCTGTTTCCCGG GA and reverse 5’ tcccgggaaacagccgccggatttggccgccacggagttctctgcagatctgccgccgga). GFP or mCherry2 with Bgl2 sites was generated by Phusion PCR (NEB) with the following primers (forward 5’ GAATTCAGATCTATGGTGAGCAAGGGCGAG and reverse 5’ gaattcagatctcttgtacagctcgtccatg) or (forward 5’ GAATTCAGATCTATGGTGAGCAAGGGCGAGGAG and reverse 5’ gaattcagatctcttgtacagctcgtccatgccg) using pEGFP (Clontech) or pCMV-mCherry2 (Clontech) as the DNA template, respectively.

#### Immunoblotting and immunoprecipitation

For western blotting, cells were washed twice in PBS, harvested in 1% NP-40 Lysis Buffer (50 mM Tris-HCl, pH 8.0, 150 mM NaCl, 1% NP-40, 0.1% SDS, 1X Protease Inhibitor Cocktail and 0.5 mM DTT) and incubated for 30 min at 4°C. Extracts were cleared by centrifugation at 14,000 × g at 4°C for 45 min and analyzed. The supernatant was removed, and protein concentrations were determined by bicinchoninic acid (BCA) assay (Pierce). Equal amounts of total protein from each sample were analyzed by SDS-PAGE on Criterion gels (Bio-Rad). SDS-PAGE samples were transferred onto Protran Nitrocellulose (GE) or Immobilon-P PVDF (Millipore) using Tris/Glycine/SDS Buffer (Bio-Rad) at 100V for one hour at 22°C. Membranes were blocked with 5% milk and 0.1% Tween-20 in Tris-buffered saline (TBS) for 1 hr at room temperature. Membranes were immunoblotted for 1 hr at 22°C using the following antibodies: rat anti-HA (1:3000; Roche), mouse anti-V5 (1:5000; Life Technologies), rabbit anti-SHH (H-160) (1:1000; Santa Cruz), rabbit Anti-GFP (1:8000, Rockland), rabbit anti-Kinesin (anti-Kif5B, 1:5000; Abcam), followed by three 5-min washes in secondary milk (primary milk diluted to 25% with TBS). Corresponding HRP-conjugated secondary antibodies (Jackson Immuno, West Grove, PA) were incubated for 1 hr at RT at a 1:5000 concentration. Blots were developed using an Odyssey Fc (Li-Cor) with ECL Prime (GE, Pittsburgh, PA).

For immunoprecipitation assays, proteins of interest were expressed in NIH3T3 cells. Cell lysates were prepared ~48 hr post-transfection using a 0.5% NP-40 Lysis Buffer (25 mM Tris-HCl, pH 7.4, 50 mM NaCl, 0.5% NP-40, 5% glycerol, 2 mM MgCl2, 1 mM EDTA, and 1X Protease Inhibitor Cocktail) and incubated for 30 minutes at 4°C with 2 units per mL of Benzonase Nuclease to degrade DNA from protein samples. Extracts were cleared by centrifugation at 14,000 × g at 4°C for 30 minutes, supernatant was collected, and protein concentration was determined by BCA assay (Pierce, Waltham, MA). Equal total protein amounts for each sample were used in co-immunoprecipitation assays and analyzed by SDS-PAGE on Criterion gels (Bio-Rad, Hercules, CA). Co-immunoprecipitation assays were performed as described (*49*) with the following modifications. Samples were pre-cleared with A/G Plus Agarose for 30 minutes with gentle rotation. Samples were centrifuged at 1000 × g for 1 minute and set up in new tubes with either anti-Mouse IgG1 control or EZview Red Anti-Flag Affinity Gel (Sigma, St. Louis, MO) (to immunoprecipitate Flag epitope-tagged proteins) for three hours at 4°C with gentle rotation. Samples were then centrifuged at 1000 × g for 1 minute and supernatant was removed. Beads were washed 3x for 5 minutes each with 0.5% NP-40 Lysis Buffer with increasing concentrations of NaCl (50 mM, 75 mM and 100 mM) with gentle rotation at room temperature. A final wash was performed with 0.5% NP-40 Lysis Buffer with 50 mM NaCl. Proteins were eluted from agarose beads with 1X SDS sample buffer (2% SDS, 4% v/v Glycerol, 40 mM Tris-HCl, pH 6.8, 0.1% Bromophenol blue) by incubating them at room temperature for 5 minutes. Samples were centrifuged at 2000 × g for 2 minutes and the eluent was transferred to a new tube. Immunoprecipitates were analyzed by western blot using the following antibodies: rabbit anti-GFP (1:8000; Rockland), rabbit anti-Flag (DDDDK) (1:3000, Abcam), rabbit anti-Shh (1:2000; SCBT), and rabbit anti-Kif5B, (1:5000; Abcam).

#### Transcriptional reporter assay

For co-culture *Gli*-reporter assays, HEK293T cells were seeded at a density of 1×10^6^ cells per 60 mm plate. The following day, pCDNA3-GFP (2µg), pCDNA3-SHH-FL-GFP-10aa linker (4µg), pCMV-mCherry2 (2µg) and pCDNA3-SHH-FL-mCherry2-10aa linker (4µg) were transfected into HEK293T cells. In a 6-well plate Light II reporter cells were seeded at a density of 0.5×10^6^ cells per well in DMEM-10% FBS complete growth media and grown overnight at 37ºC, 5% CO_2_. The following day, transfected HEK293T cells underwent trypsinization and were seeded into the Light II wells at a density of 0.5×10^6^ cells per well. Cells were allowed to recover for 4 hrs at 37ºC, 5% CO_2_. Media was removed from cells, washed twice with PBS and once with DMEM serum-free complete media (phenol red free). DMEM serum-free complete media was added back to each well and allowed to incubate for 2 hrs. Washing was carried out over 6 hrs repeating the above wash steps. After 6 hr, 3 mL of DMEM Serum-free Complete Media was added to each well and the cells were incubated for ~36 hr. Reporter assays were carried out according to Dual Luciferase Reporter Assay Kit instructions (Promega). Experiments were repeated three times in triplicate.

#### Electron Microscopy

NIH3T3 cells expressing SHH-mCherry were seeded at 60% confluency into 8 well, Permanox slide, 0.8 cm^2^/well (Nunc Lab-Tek II) and, for pre-embedding immunolabeling, were fixed in a 0.5% glutaraldehyde with 4% PFA fixative in 0.1 M phosphate buffer. Prior to labeling with primary antibody, samples were washed with buffer and excess aldehyde groups neutralized with glycine. Samples were blocked with 1% BSA in 10mM PBS (BSA/PBS) then a blocking solution matched to the species of the secondary antibody (Aurion, Wageningen, The Netherlands) in PBS. Samples were incubated with chicken anti-mCherry (1:1000, abcam) diluted in BSA/PBS overnight at 4°C. Following primary antibody incubation, samples were washed in BSA/PBS then incubated with a streptavidin conjugated secondary antibody. Samples were rinsed with PBS and incubated with biotinylated nanogold (Nanoprobes, Yaphank, NY) then washed with PBS and fixed in 1% glutaraldehyde (Electron Microscopy Sciences (EMS), Hatfield, PA). Following fixation, samples were successively rinsed in distilled water and 0.2 M citrate buffer then incubated with HQ Silver Enhancement reagent (Nanoprobes) prepared per manufacturer instructions. Enhancement reaction was halted by rinsing with distilled water. Samples were contrasted successively with 1% osmium tetroxide (EMS) and 1% uranyl acetate (EMS) in water with water washes between contrasting steps. Samples were then dehydrated in an ascending series of alcohols, infiltrated with EmBed 812 (EMS) and polymerized at 80°C overnight. Samples were sectioned on a Leica ultramicrotome (Wetzlar, Austria) at 70nm and examined in a Tecnai G² F20-TWIN transmission electron microscope. Images were recorded using an AMT side mount camera system. Unless specified, all chemical and reagents were from Sigma (St. Louis, MO).

## Acknowledgements

We thank members of the Ogden lab for thoughtful discussion during the course of this work and for comments on the manuscript. We thank Ben Allen, Adrian Salic and Robert Krauss for BOC, CDON and GAS1 expression vectors and knockout cell lines.

## Funding

This work was supported by National Institute of General Medical Sciences grants R35GM122546 (SKO), R01GM134531 (REC) and by ALSAC of St. Jude Children’s Research Hospital. The CTI is supported by the Cancer Center Support Grant, NCI P30 CA021765.

## Authors contributions

**Conceptualization** ETH and SKO

**Methodology** ETH, DPS, MD, JT, MM, CGR and SKO

**Software** ETH and JT

**Validation** ETH, DPS, MD, BW and AS

**Formal analysis** ETH, AS and MM

**Investigation** ETH, DPS, MD and BW

**Resources** JT, REC, MM, CGR and SKO

**Writing** ETH and SKO

**Visualization** ETH

**Supervision** SKO

**Funding acquisition** REC and SKO

## Competing interests

Authors report no conflicts of interest.

## List of Supplementary Materials

Materials and Methods

Figure S1-S9

Movie S1-7

## Notes

### Competing Interest Statement

The authors have declared no competing interest.

