## Supplemental Material for "Myosin 10 and a Cytoneme-Localized Ligand Complex Promote Morphogen Transport"

**This PDF file includes:**

Figs. S1 to S9
Captions for Movies S1 to S7

**Other Supplementary Materials for this manuscript include the following:**

Movies S1 to S7

### Supplemental Figures

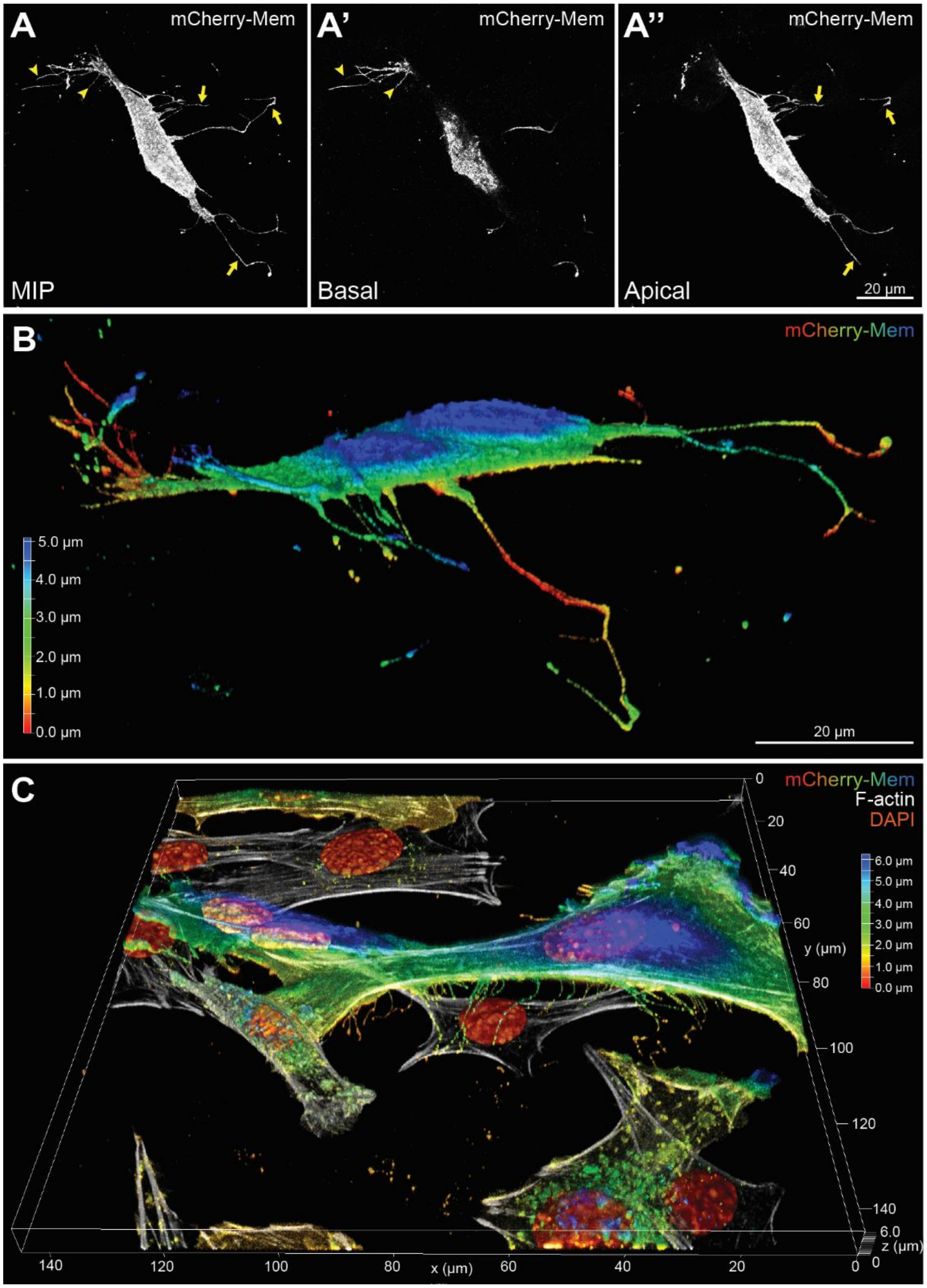


**Supplemental Figure 1: Cytonemes originate from cell membrane not in contact with the growth substrate.** (**A-B**) A MEM-fixed NIH3T3 cell expressing mCherry-Mem and SHH is shown in support of **Fig. 1A, A'**. (**A**) Maximum intensity projection (MIP) of the cell identifies both coverslip-anchored membrane extensions (arrowheads) and cytonemes (arrows). (**A'**) Cell membrane protrusions adjacent to the culture coverslip (basal) are shorter. Cytoneme sections are detectable making contact with the coverslip, but do not originate from membrane in contact with the coverslip. (**A''**) Projection of cell membrane not in contact with the coverslip (apical) shows cytoneme initiation. (**B**) Lateral 3D render with Z-axis depth shading of mCherry-Mem. The depth scale is shown. (**C**) A 3D render of a SHH and mCherry-Mem (Z-depth shaded) expressing MEF showing cytonemes and filopodia in support of **Fig.1 B,B'**.

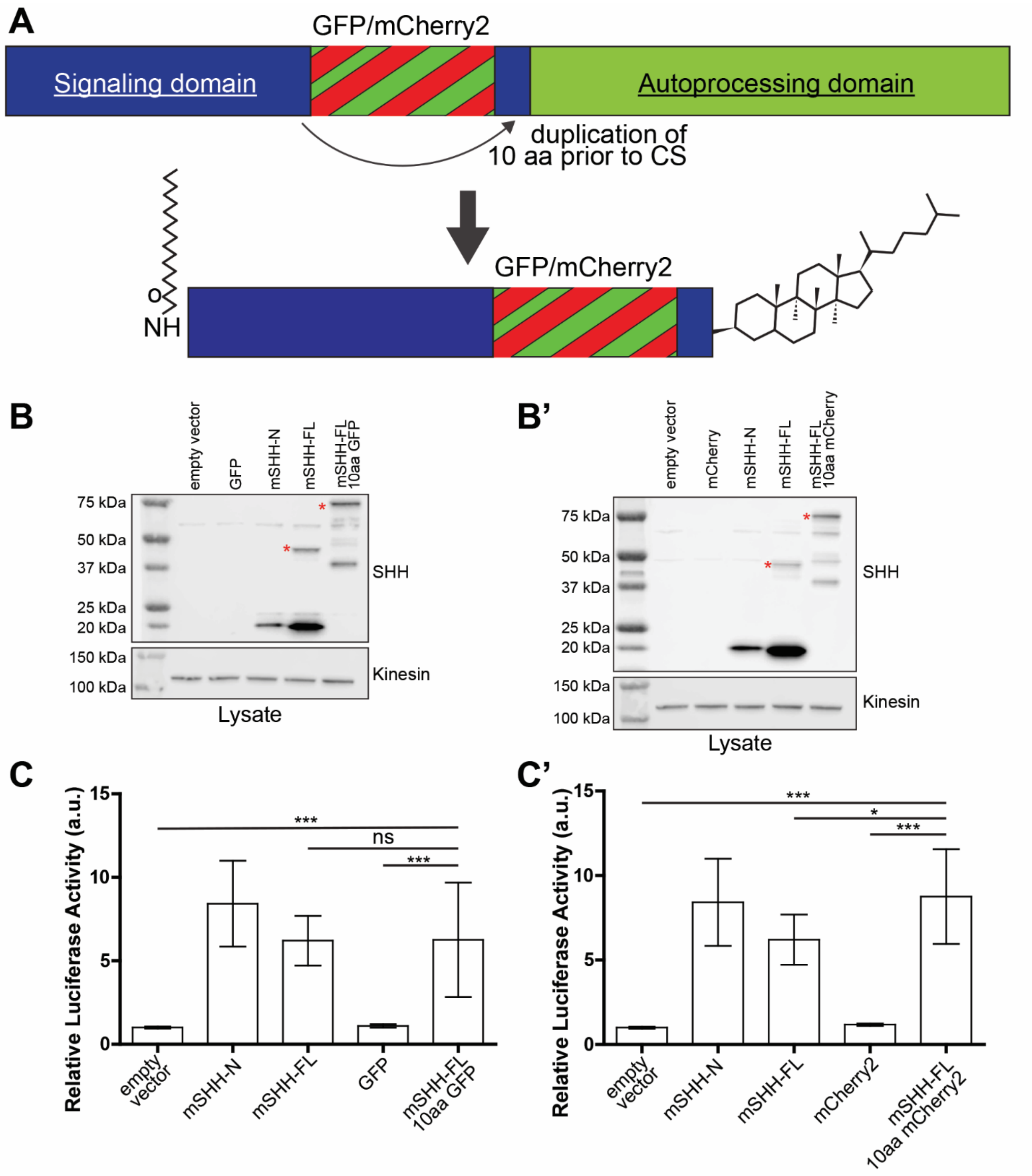


**Supplemental Figure 2: Generation and validation of mouse (m)SHH-GFP and SHH-mCherry.** (**A**) GFP or mCherry2 were inserted into the SHH signaling domain immediately prior to the autoprocessing cleavage site (CS). The 10 amino acids prior to the cleavage site (CS) were duplicated carboxyl-terminal to the insert to ensure normal processing. (**B,B'**) Immunoblots of lysates from NIH3T3 cells expressing (**B**) SHH-GFP or (**B’**) SHH-mCherry are shown compared to SHH-FL and SHH-N. Red asterisks indicate unprocessed SHH proteins. (**C,C’**) SHH responsive LightII cells were co-cultured with HEK293T cells expressing the indicated SHH-fluorescent fusion proteins and assayed for ligand-induced reporter gene induction. Reporter assays were performed in triplicate, with 3 biological replicates. All data are presented as mean ± SD. ns = not significant, * $p<0.05$, *** $p<0.001$.

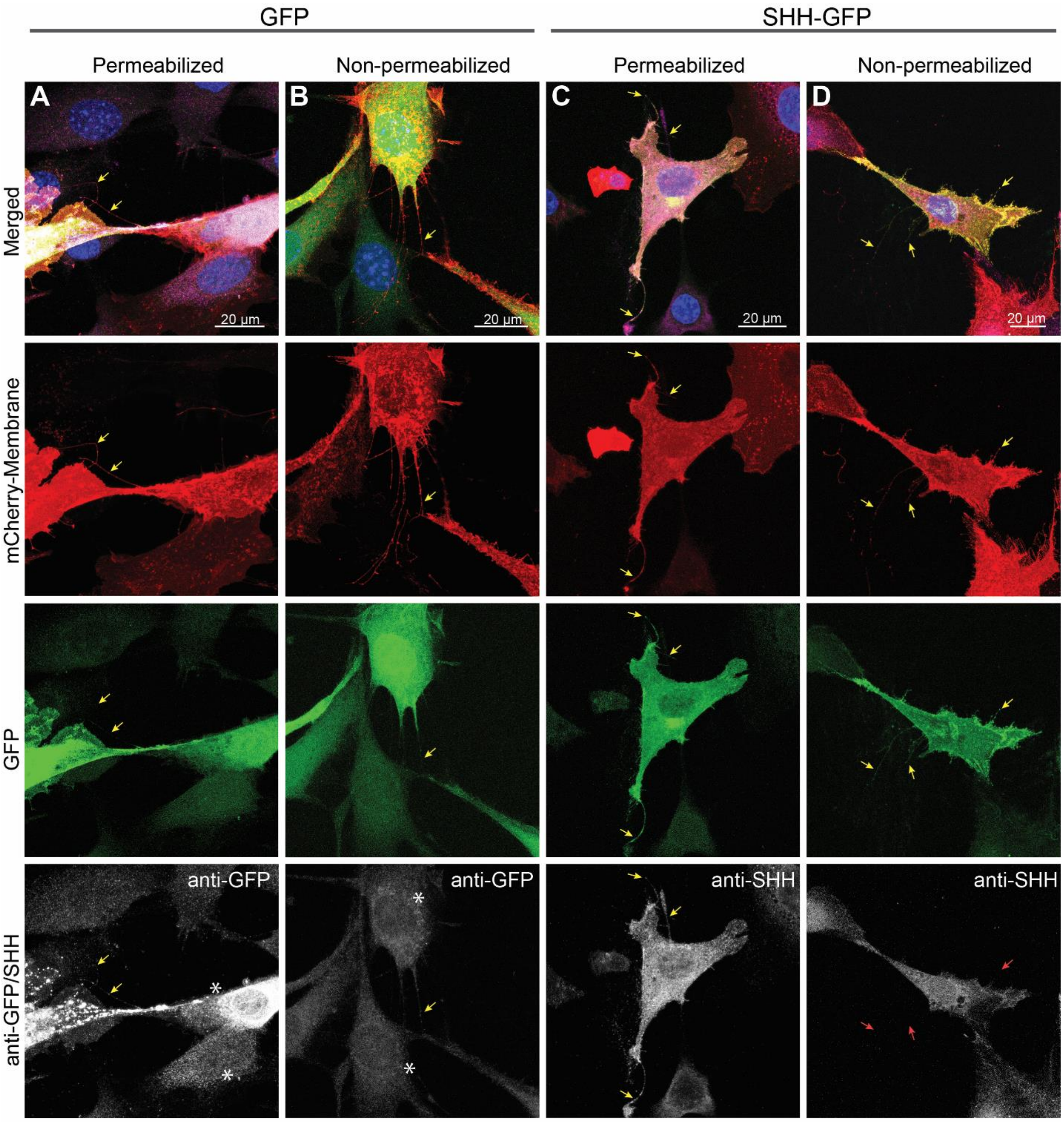


**Supplemental Figure 3: Control experiments for non-permeabilization staining of SHH and GFP.** (**A,B**) Cytoplasmic GFP and mCherry-Mem transfected NIH3T3 cells were immuno-stained for GFP under, (**A**) permeabilized or, (**B**) non-permeabilized conditions. GFP is detected by anti-GFP in permeabilized cells, but not in non-permeabilized cells (asterisks). (**C,D**) SHH-GFP and mCherry-Mem transfected NIH3T3 cells were immuno-stained using anti-SHH under, (**C**) permeabilized or, (**D**) non-permeabilized conditions. SHH-GFP signal is evident in cytonemes in both conditions (yellow arrows), but not evident in non-permeabilized cells when probes with anti-SHH (red arrows).

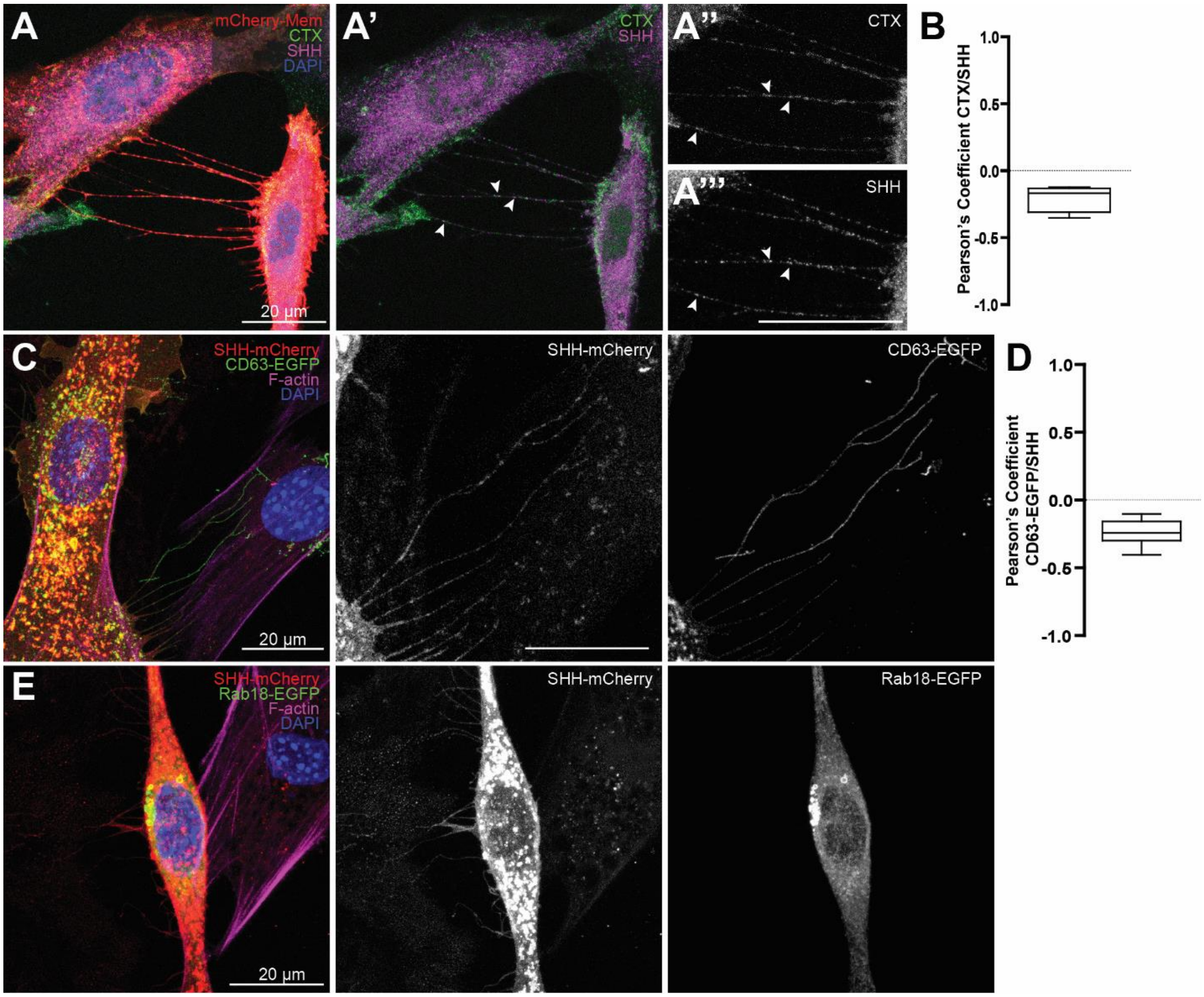


**Supplemental Figure 4: SHH does not colocalize with lipid rafts, CD63 or Rab18 in cytonemes.** (**A-A'''**) NIH3T3 cells expressing SHH (magenta) and mCherry-Mem (red), were incubated with lipid raft marker CTX (green) and analyzed by confocal microscopy. Arrowheads indicate rare colocalization events between SHH and CTX along cytonemes (**A'** shows CTX and SHH, **A'',A'''** shows single channels zoomed in). (**B**) Box plot with min/max whiskers of Pearson's coefficients values of colocalization between SHH and CTX in cytonemes. (**C-E**) Cytonemes of NIH3T3 cells expressing SHH-mCherry and CD63-EGFP (**C**) or Rab18-EGFP (**E**) analyzed by confocal microscopy do not show significant co-localization. Rab18-EGFP does not enter cytonemes. (**D**) Box plot with min/max whiskers of Pearson's coefficients values of colocalization between SHH-mCherry and CD63-EGFP in cytonemes.

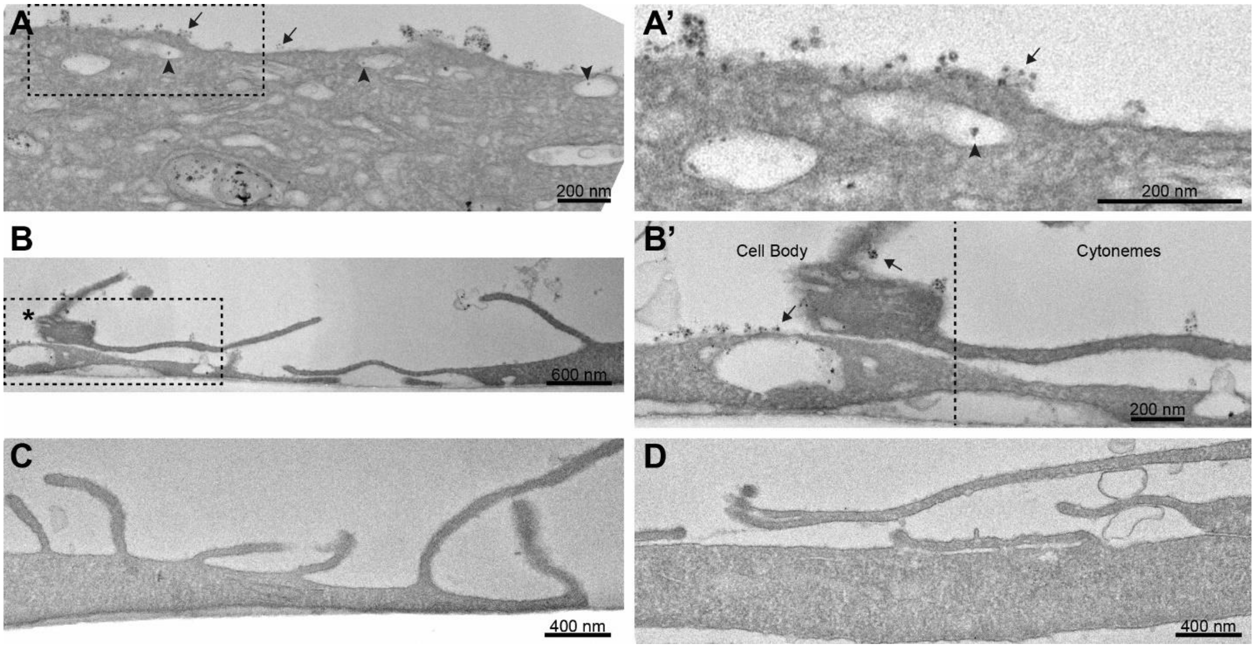


**Supplemental Figure 5: Immuno-TEM of SHH-mCherry in NIH3T3 cell cytonemes.** (**A,A’**) Anti-mCherry immunolabelling detects SHH-mCherry in interior vesicles (arrowheads) and along the plasma membrane of producing cells (arrows, **A’** zoom). Plasma membrane localization is similar to what was observed in nonpermeabilized immunofluorescent stains (**Fig. 2A**). (**B-B’**) SHH-mCherry producing cell (asterisk, left) and receiving cells (right). (**B’**, zoom) Surface SHH-mCherry clusters are present along the main cell body (arrows), but absent along cellular protrusions. (**C**) Sections of SHH-mCherry-expressing NIH3T3 cells not exposed to mCherry primary antibody were incubated with streptavidin conjugated secondary antibody followed by silver enhancement. No artifact signals were detected. (**D**) A section from an NIH3T3 cell not expressing SHH-mCherry was incubated with anti-mCherry prior to secondary antibody incubation and silver enhancement. No nonspecific labelling was detected.

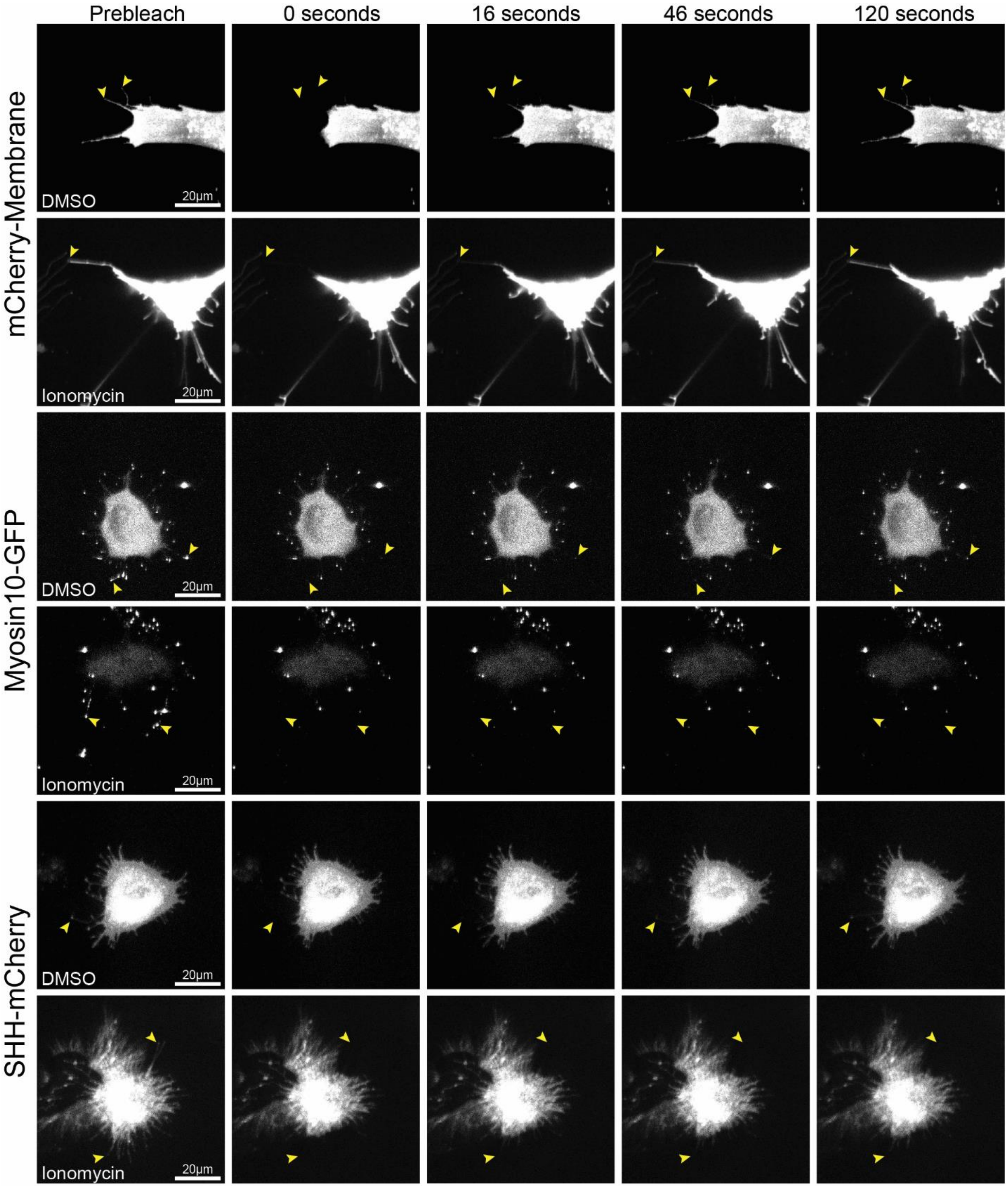


**Supplemental Figure 6: FRAP of NIH3T3 cell cytonemes treated with DMSO (control) or ionomycin.** Cytoneme tips are indicated by yellow arrowheads. Ionomycin does not impair mCherry-Membrane recovery along the cytoneme. MYO10-GFP and SHH-mCherry fail to recover to cytoneme tips by 120 seconds post-photobleaching in treated cells.

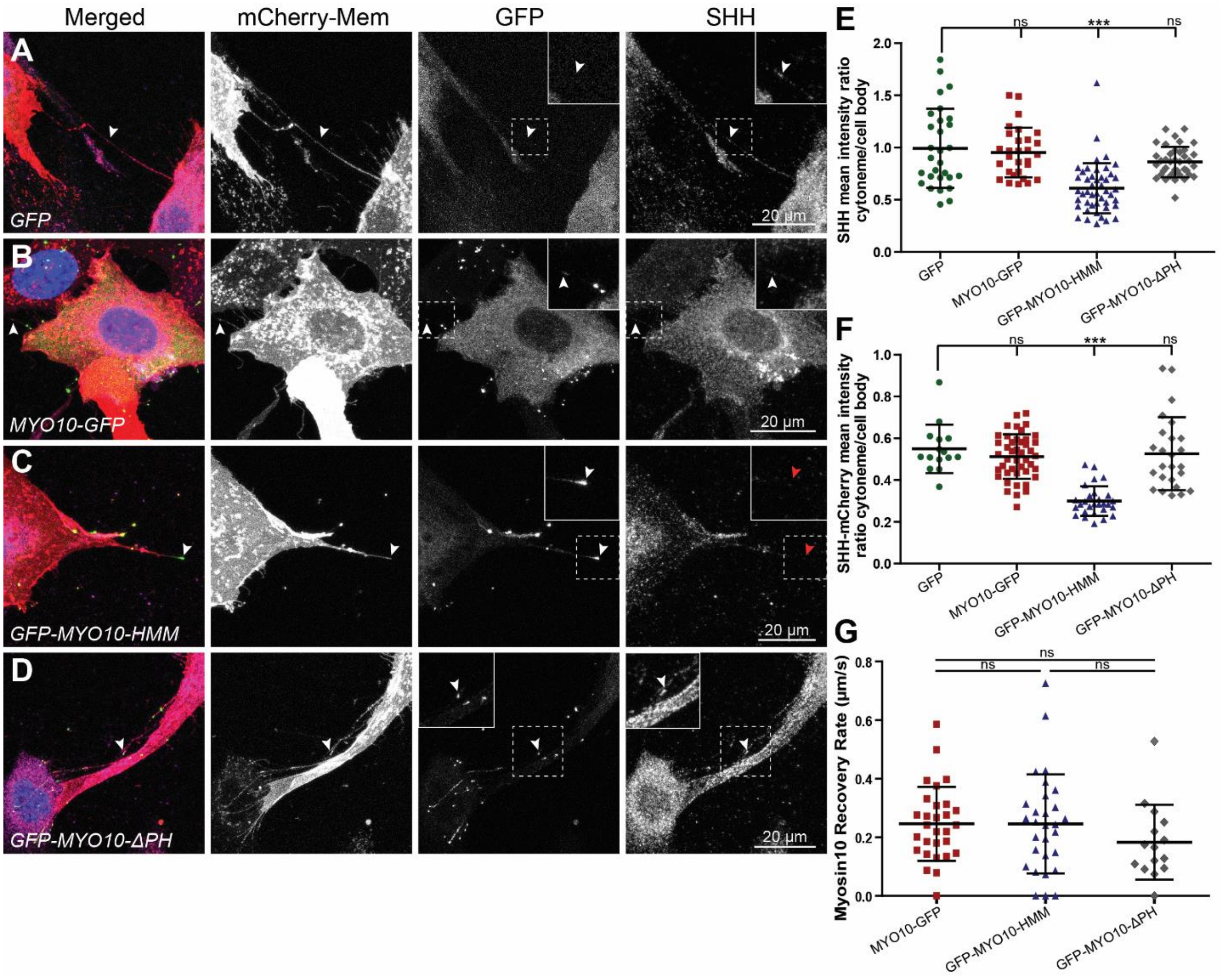


**Supplemental Figure 7: MYO10 influences SHH transport in cytonemes.** (**A-D**) NIH3T3 cells expressing mCherry-Mem and SHH with either (**A**) GFP, (**B**) MYO10-GFP, (**C**) GFP-MYO10-HMM, or (**D**) GFP-MYO10-ΔPH are shown. Arrowheads indicate cytonemes. Red arrowheads mark a MYO10-HMM positive cytoneme that fails to accumulate SHH. (**E,F**) Scatter plots of mean (**E**) SHH, or (**F**) SHH-mCherry fluorescent ratios of cytoneme:cell body in NIH3T3 cells when co-expressing GFP or the indicated MYO10 constructs. (**G**) Scatter plots of recovery rates of MYO10 constructs to cytoneme tips. All data are presented as mean ± SD. ns = not significant, *** $p<0.001$.

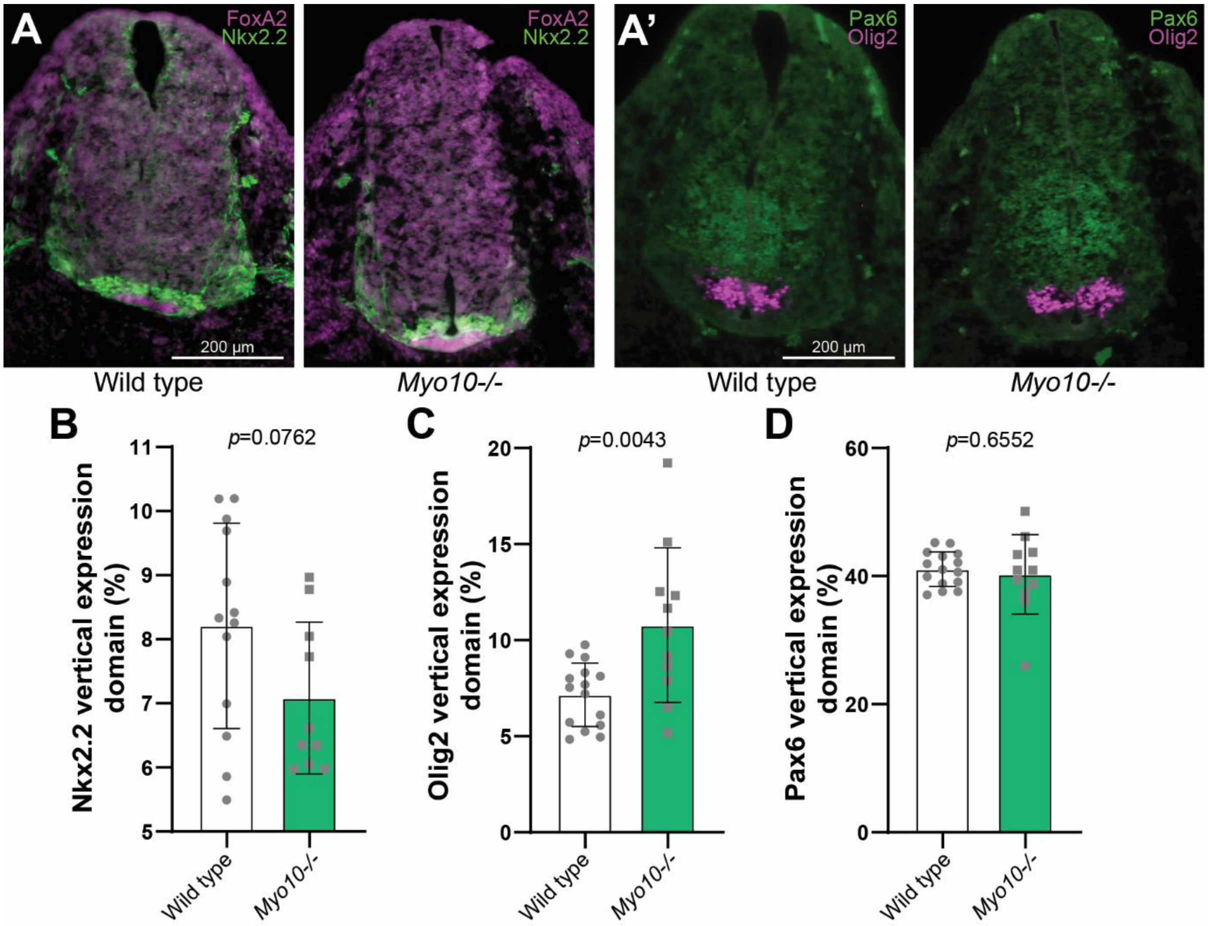


**Supplemental Figure 8: Anterior *Myo10-/-* neural tube progenitor domains are altered.**

(**A,A'**) Cardiac-level neural tube sections of wild type and *Myo10*$^{m1J/m1J}$ E10.5 embryos at the were immuno-stained using (**A**) anti-FOXA2 (magenta) and anti-NKX2.2 (green), or (**A'**) anti-PAX6 (green) and anti-OLIG2 (magenta). (**B-D**) Relative expression domains of (**B**) *Nkx2.2*, (**C**) *Olig2*, and (**D**) *Pax6* in wild type and *Myo10*$^{m1J/m1J}$ mice indicate expansion of *Olig2* and compression of *Nkx2.2*. Each dot represents an individual section from the representative animal (n=3 embryos per genotype).

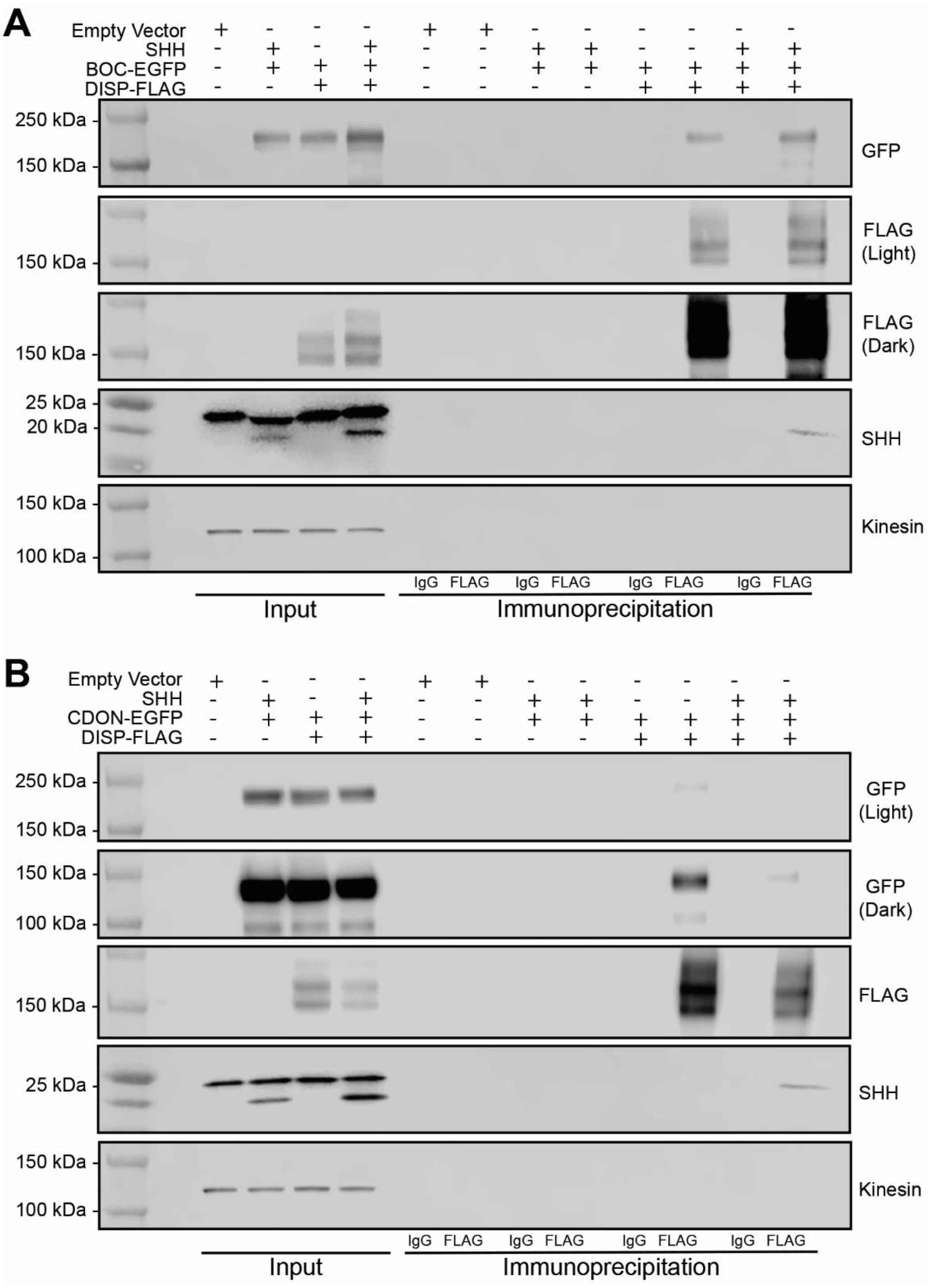


**Supplemental Figure 9: Western blots in support of Figure 4J-K.** Immunoblots of lysates and immunoprecipitates from NIH3T3 cells expressing DISP-FLAG and BOC-EGFP (**A**) or CDON-EGFP (**B**) with or without SHH. Anti-FLAG immunoprecipitation with input (10%).

**Movies**

**Supplemental Movie 1: Cytonemes are dynamic cellular extensions that contain MYO10.** Two adjacent NIH3T3 cells expressing SHH, mCherry-Mem (magenta), and MYO10-EGFP (green) are shown as a maximum intensity projection of 10 Z-sections spanning 4.5 µm, imaged at 4.1 seconds/frame over 18 minutes. Cytonemes are distinguished from other cellular extension by active growth and accumulation of MYO10-EGFP at the tips. Time stamp indicates minutes:seconds.

**Supplemental Movie 2: Dynamic cytonemes move in 3 dimensions.** Lateral projection of a cell edge from Movie S1, imaged as described in Supplemental Movie 1. Active cytoneme extensions frequently traverse through the media, eventually dropping onto the culture surface. Time stamp minutes:seconds.

**Supplemental Movie 3: Cytonemes exhibit transient interactions and stable connections.** NIH3T3 cells expressing SHH, mCherry-Mem (magenta), and MYO10-EGFP (green) are shown as a maximum intensity projection of 5 Z-sections spanning 2 µm, imaged at 2.05 seconds/frame over 9 minutes. Cytonemes transiently scan membrane of a neighboring cell (upper right) and form stable connections with cytonemes from adjacent cells (center). MYO10-GFP moves along cytonemes in puncta and enriches at cytoneme contact points. Time stamp indicates minutes:seconds.

**Supplemental Movie 4: NIH3T3 R-GECO positive cells in contact with GFP expressing cells.** GFP-expressing NIH3T3 cells (green) are shown in contact with R-GECO reporter cells (orange intensity spectrum) and presented as a maximum intensity projection of 4 Z-sections spanning 3 µm, imaged at 1.7 seconds/frame over 15 minutes. Time stamp indicaes hours:minutes:seconds.

**Supplemental Movie 5: NIH3T3 R-GECO positive cells in contact with cytonemes from SHH-GFP expressing cell.** Cytonemes from SHH-GFP expressing cells (green) are shown contacting R-GECO reporter cells (orange intensity spectrum), shown as a maximum intensity projection of 4 Z-sections spanning 2.5 µm, imaged at 1.7 seconds/frame over 15 minutes. Time stamp indicates hours:minutes:seconds.

**Supplemental Movie 6: FRAP of NIH3T3 cytonemes expressing SHH-mCherry and MYO10-GFP treated with DMSO.** SHH-mCherry (white) and MYO10-GFP (green) recover to the cytoneme tip within 120 seconds post photobleaching. Cell was imaged at 2 frames per second from a single focal plane for 5 seconds prior to photobleaching, following ~130 seconds of recovery.

**Supplemental Movie 7: FRAP of NIH3T3 cytonemes expressing SHH-mCherry and MYO10-GFP treated with ionomycin dissolved in DMSO.** SHH-mCherry (white) and MYO10-GFP (green) do not recover to the cytoneme tip post photobleaching. Recovery of SHH-mCherry occurs along the base of some cytonemes but fails to reach the tip. Live images were acquired as described for Supplemental Movie 6.
